# An ERF transcription factor PTI5, a novel regulator of endophyte community maintenance in potato

**DOI:** 10.1101/2025.04.24.650297

**Authors:** Tjaša Lukan, Barbara Kraigher, Karmen Pogačar, Katja Stare, Teja Grubar Kovačič, Maja Zagorščak, Marko Petek, Polonca Stefanic, Anže Vozelj, Valentina Levak, Tjaša Mahkovec Povalej, Juan M. García, Maria J. Pozo, Ines Mandić-Mulec, Kristina Gruden

**Author notes:** These authors share last authorship (Ines Mandić-Mulec, Kristina Gruden). corresponding authors Kristina Gruden.

## Abstract

We have recently identified ERF transcription factor PTI5 as a susceptibility factor, negatively regulating immune response to diverse pathogens. Here we investigated the processes involved in colonisation of potato with beneficial organisms. RNAseq showed that at the time of *Bacillus subtilis* biofilm establishment, immune responses in interacting roots were attenuated, and complex transcriptional network was triggered, with ethylene signalling being a central module and PTI5 strongly induced. Interestingly, the response is intensified if plants are inoculated by two antagonistic *B. subtilis* strains. While PTI5 is not involved in the establishment of biofilm on roots, we show that bacterial abundance increases in PTI5-silenced plants. Remarkably, root colonization by the arbuscular mycorrhizal fungus *Rhizophagus irregularis* was also higher in the PTI5-silenced plants. PTI5 is thus involved both in blocking defence against harmful and blocking colonisation with beneficial microbes. Such mechanistic understanding of plant–microbe interaction paves the way for sustainable crop management.

## INTRODUCTION

Plant-microbe and microbe–microbe interactions in the rhizosphere determine plant health and productivity.^1^ Beneficial microorganisms are common in rhizosphere soils. Some can endophytically colonize plants, frequently improving plant nutrition and plant growth, and protect plants from disease and abiotic stresses through a wide variety of mechanisms.^2,3^ They can be efficient and environmentally friendly alternatives to chemical pesticides and fertilizers, due to their multiple benefits for the plants and their long shelf life, which is comparable with that of agrochemicals.^4^ However, the current limitation for a wider adoption of microbial inoculants in agriculture is reproducibility of the results under field conditions.^5^ Additional knowledge, such as a deeper understanding of the mechanisms involved in plant colonization and maintenance of the plant microbiome is required to improve the use of microbes as plant protectants.^6^

The plant immune system is central in shaping the plant response to beneficial and pathogenic microorganisms, regulating the extension and successful colonization of plant tissues by mutualistic organisms and preventing the invasion of the parasitic ones. Recognition of pathogens leads to the activation of interconnected plant hormonal signaling pathways mediating downstream transcriptional reprogramming resulting in effective defense responses.^7,8^ When plant interacts with beneficial microbes, microbe triggered immunity (MTI) is activated similarly as in interaction with pathogens.^9^ Beneficial microbes developed several strategies to overcome this immune response and colonize the plant. One option to avoid effective recognition is by modifying the microbe-associated-pattern, recognized by immunity, for example flagellin. Alternatively, the microbes develop tolerance to toxic compounds produced, e.g. reactive oxidative species (ROS) or they actively suppress MTI.^10^ They can also modify metabolism of plant to allow for easier penetration, for example suppress synthesis of suberin.^11^

Successful colonization by beneficial microbes can induce resistance to pathogenic bacteria and herbivores. Mechanisms of induced resistance (IR) are not well understood, but it has been shown to involve jasmonic acid (JA), salicylic acid (SA) and ethylene signaling pathways.^11, 12, 13^ Several individual transcription factors have been shown also to be important for the establishment of the IR state. As described for bacteria, root colonization by beneficial fungi can also boost plant immunity. For example, root colonization by arbuscular mycorrhizal fungi is tightly regulated and the modulation of immune responses during the symbiosis establishment can also leads to induced resistance in systemic tissues.^15^ Therefore, understanding the fine-tuned regulation of plant immune responses during colonization by beneficials is essential for optimizing bioinoculant-based strategies in agriculture, allowing their establishment and promoting their benefits in plant health.

In the case of bacteria, social behavior has been proven to be crucial for the success in colonizing plant tissues. Bacteria often exist in multicellular groups of cells called biofilms,^16^ where they engage in competition for resources but also in cooperative (synergistic) interactions that enhance community productivity.^17^ These social engagements are most often mediated through secreted molecules that are believed to have a profound impact on bacterial diversity, spatial organization, and ecological functions.^16,18^ *B. subtilis* species are capable of different forms of social interactions besides biofilm formation, such as quorum sensing (QS), group motility (swarming), and kin discrimination.^19^ Non-kin *B. subtilis* strains form a striking boundary between approaching swarms,^20,21^ exclude a non-kin competitor from a common swarm,^22^ and do not mix on *Arabidopsis thaliana* roots, while kin strains merge on solid medium and cohabit root surfaces.^20^ *B. subtilis* strains also produce a range of antibiotics, surfactants, secondary metabolites, and bioactive volatiles expected to modulate social interactions between bacteria and with the plant.^20,21,23,24^ *B. subtilis* QS system comprises the signalling peptide ComX, which positively regulates the synthesis of surfactin (encoded by Srf operon), one of the most potent surfactants and a lipopeptide antibiotic.^19,25^ Surfactin acts as a signalling molecule during biofilm formation, promotor of horizontal gene transfer and as an inducer of physiological changes in plants during colonization with bacteria.^25,26,27,28,29^ Despite evidence for *in situ* QS dependent expression of surfactin, the link between QS and surfactin on plant roots and how QS affects plant colonisation and plant immune response is less understood.

As potato, the third most important crop in the world, is exceptionally sensitive to a wide range of environmental stresses,^31^ studying its interaction with beneficial microbes is crucial to ensure an efficient environmentally friendly system of plant protection based on modulation of its microbiome. The aim of this study was to investigate mechanisms involved in the interaction between potato and beneficial microbes. Different *B. subtilis* strains and mutants were used to investigate the principles of bacterial social interactions and their interaction with the plant. We identified transcription factor PTI5 as a potential regulatory hub for *B. subtilis* endophytic colonization and further investigated its role in the plant–bacteria interaction using transgenic plants with silenced PTI5. We here show that *B. subtilis* abundance is increased in systemic tissue of plants with silenced PTI5, suggesting that PTI5 maintains optimal *B. subtilis* abundance in colonized plants. Interestingly, a similar function was confirmed also for the establishment of the mutualistic symbiosis between potato and arbuscular mycorrhizal fungi.

## RESULTS

### Stable *B. subtilis* biofilm formation on potato roots coincides with first transcriptional responses in plant roots

We here studied the response of potato to colonization by two different *B. subtilis* strains (PS-216 and PS-218). Both formed biofilm on potato roots (Figure S1). As our aim was to capture both initial events in biofilm establishment as well as long-term effects of potato–*B. subtilis* interaction, we first determined timing of biofilm establishment on potato roots and transcriptional responses in the plant. We selected eight genes known to be involved in potato immune response as potential indicators of potato MTI to *B. subtilis* colonization^32,33^ and followed their expression in potato roots and shoot tissue. The genes showed diverse expression profiles, which were for some genes tissue specific (Figure S2, Table S1). Two genes, an ERF transcription factor PTI5, which requires an active ethylene pathway for the activation and mediates crosstalk of ethylene and SA pathways^33^ and a gene involved in JA synthesis, 13LOX,^32^ were reproducibly upregulated in roots and shoots 24 hours post-bacterial inoculation (Figure 1, Figure S2, Table S1). The results were similar for both tested *B. subtilis* strains (Exp1 and Exp2 in Figure S2, Table S1).

**Figure 1:**
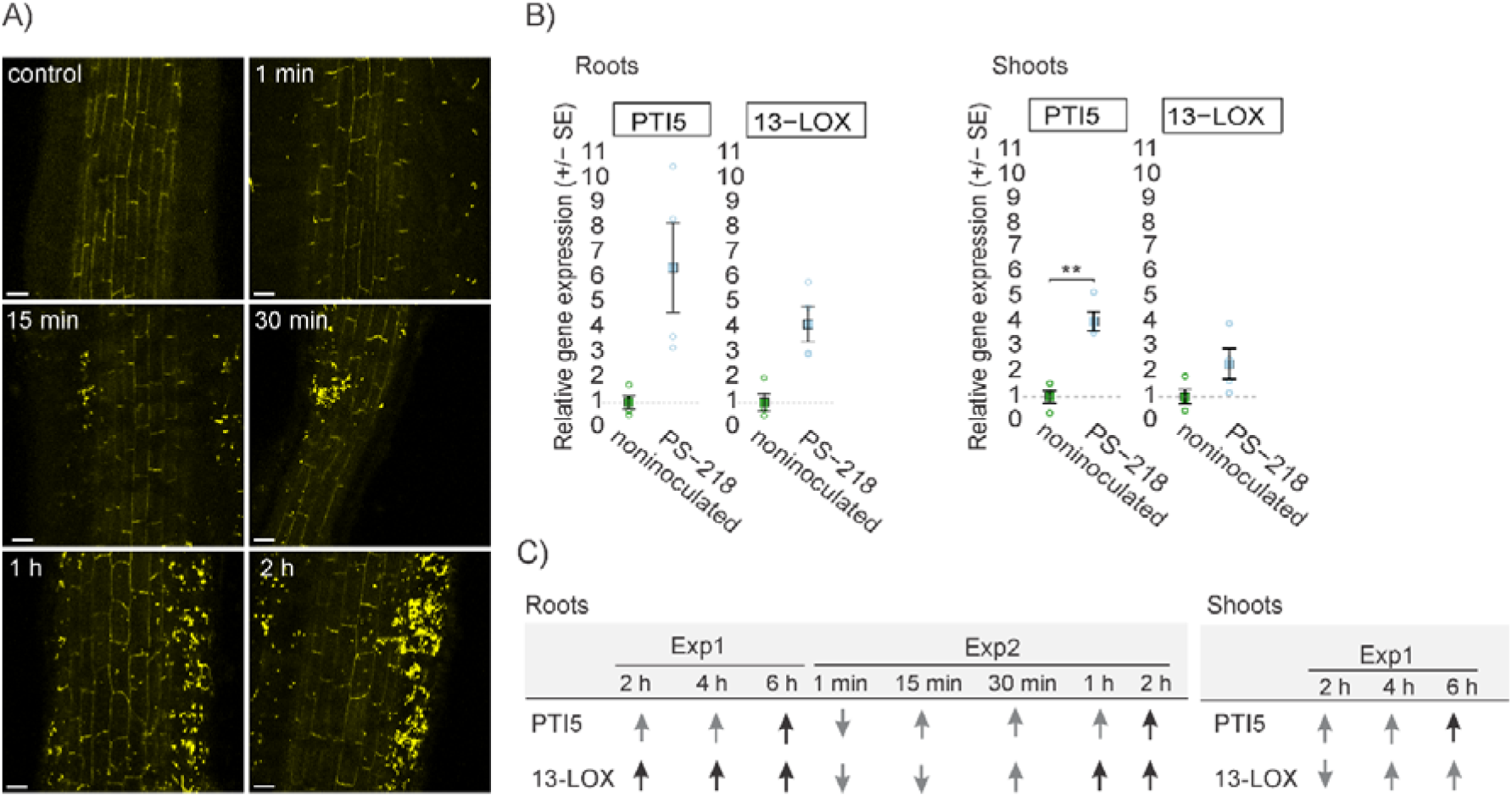
Dynamics of biofilm formation and plant responses of potato to *B. subtilis*. A) Biofilm formation (YFP fluorescence) was followed on potato roots after incubation with *B. subtilis* PS-218 YFP for a specified time. Scale is 50 µm. B) Potato response to bacteria colonization is robustly followed by expression of PTI5 and 13-LOX. Relative expression of PTI5 and 13-LOX in roots and shoots of *B. subtilis* PS-218 YFP inoculated and non-inoculated plants, after overnight incubation. Relative gene expression was scaled to the average gene expression of non-inoculated group. Welch’s t-test was used to determine differences between treatments. Individual measurements (circles), mean (squares) and standard error are shown. Asterisks (*) denote a statistically significant difference (p-value < 0.05). Data from all experiments are given in Table S1. C) Dynamics of PTI5 and 13-LOX expression (up- or downregulation) in roots and shoots of potato plants after inoculation with high *B. subtilis* strain PS-218 culture density (10^7^ CFU/mL) and incubation for a specified time. Data of all experiments are available in Table S1. Up- or downregulation was determined by comparing expression in inoculated plants with expression in non-inoculated plants. ↑: upregulation, ↓: downregulation, ↑: statistically significant upregulation determined by Welch’s t-test, ↓: statistically significant downregulation determined by Welch’s t-test. Exp: experiment.

To study the dynamics of biofilm formation, we set up an experiment with higher density bacteria inoculum (10^7^ CFU/mL). In such set-up, biofilm reproducibly began forming after the first 15 minutes of incubation, and it was consistently present over the entire root surface after a 2-hour incubation period (Figure 1). Interestingly, the induction of PTI5 and 13-LOX genes was also detected when the biofilm was formed uniformly on the root (Figure 1, Figure S2, Table S1).

### Complex regulatory network is triggered when potato is colonized with *B. subtilis*

To investigate plant response to *B. subtilis* colonization in more detail, we next performed RNA-seq analysis of colonized roots and systemic shoot tissue. Our special interest was in the starting point of colonization, as we intended to capture initial *B. subtilis* recognition related responses in the plant. Therefore, roots were analyzed at the time of biofilm formation (2 hours post-inoculation (hpi)) and 24 hours later (26 hpi). The response of the plant was stronger at later time points (Figure 2A, Figure S3). Moreover, shoots were analyzed at this later time to explore potential activation of MTI and/or IR. For both root and shoot tissue, we identified a large overlap in differentially expressed genes for plants inoculated with two different strains (Figure 2A, Figure S3).

**Figure 2:**
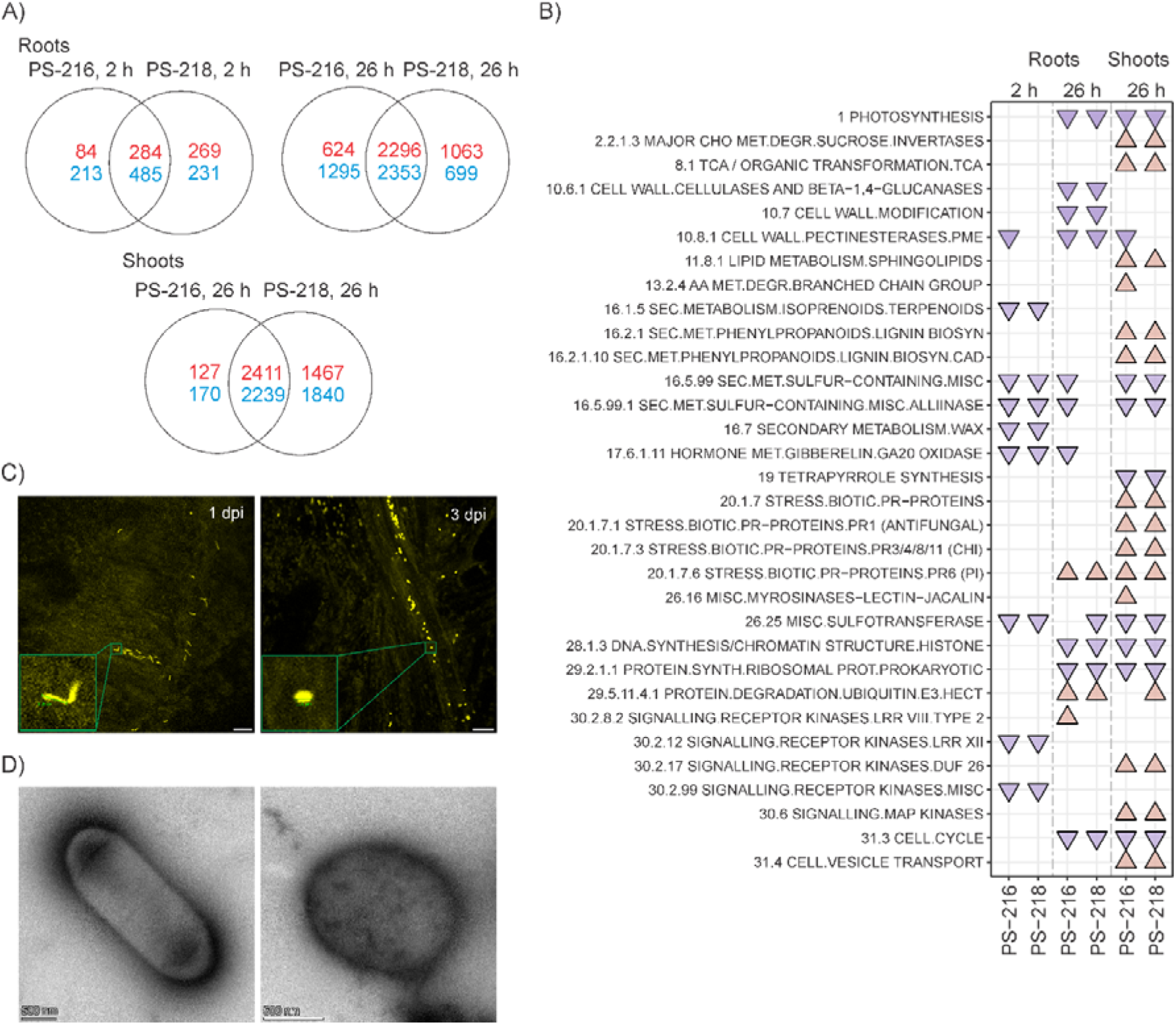
Potato adjusts to colonization with *B. subtilis* by repression of immune signaling and cell wall synthesis. A) Number of genes, regulated by *B. subtilis* in plant roots and shoots after 2-h and 26-h incubation with PS-216 YFP and PS-218 mKate2. Blue: downregulated genes, red: upregulated genes. B) Visualization of results in the context of biological pathways. For each comparison between *B. subtilis*-inoculated and non-inoculated plants, gene sets enriched for up- or downregulated genes (q-value < 0.1) are shown and designated as red triangles or blue triangles for shoots and roots at different time points. MapMan ontology BINs were used to generate gene sets. Sample group comparisons of inoculated to non-inoculated (in columns) are designated with bacterial strain names used for inoculation. PS-216: PS-216 YFP, PS-218: PS-218 mKate2. See Table S2 for all results. C) *B. subtilis* morphology is changed after internalization in plants. Left: *B. subtilis* morphology 1 dpi (day post-inoculation). Right: *B. subtilis* morphology 3 dpi. See Figure S4 for more data. Scale is 25 µm. D) Electron micrograph of free-living (left) and *in planta* living *B. subtilis* PS-216 (right). Scale is 500 nm.

Many processes related to primary line of defense were downregulated in roots at the time of biofilm formation, including diverse immune receptors, genes involved in cell wall modifications and in synthesis of terpenoids and waxes (Figure 2B). Interestingly, among the genes that were upregulated in roots already at the time of biofilm formation were in majority genes coding for transcription factors, hormonal synthesis, signalling components and components of protein degradation. Several ERF transcription factors were upregulated besides the already identified PTI5, as well as many MYBs, WRKYs, bZIP and NAC transcription factors. We also identified several lipoxygenases upregulated (JA synthesis), but also genes involved in ethylene and gibberellin synthesis as well as in Ca^2+^ signalling (Table S3). The response of most of these genes was intensified in later time point in both roots and shoots (Table S3).

Remarkably, more induced gene sets were detected in shoots. The processes activated in shoots resemble IR as studied so far in other plants. Receptor kinases were upregulated as were MAP kinases and executors of plant defense, such as beta-glucanases, alpha amylase inhibitors, PR1 proteins and PR5 proteins (Figure 2B, Table S2, Table S3). Among the genes associated with transcriptional regulation, NPR1 was also specifically upregulated in shoots only.

In addition, we also followed the extent of bacterial internalization within the plant. We detected bacteria in stem already 1 day post-inoculation (dpi) (Figure S4). Interestingly, while plant responds to *B. subtilis* colonization, also *B. subtilis* adapts once internalized as its morphology starts to change to more rounded form as early as 2 dpi (Figure 2C, Figure S4). Also using electron microscopy, we can see that most free-living bacteria are multiplying, while no multiplication was detected in *in planta* living bacteria (Figure 2D, Figure S5).

### Biofilm formation and plant responses are attenuated in potato roots inoculated with *B. subtilis* quorum sensing and surfactin-deficient mutants

To better understand the mechanisms that trigger plant responses, we inspected the role of the *B. subtilis* ComX peptide mediated QS and surfactin production in potato–*B. subtilis* interaction. We used two PS-216 mutant strains, one with deletion of the *comQXP* gene cluster, coding for the genes involved in QS, which also regulate surfactin production (PS-216 Δc*omQXP*),^25^ and the other strain with a mutation in the *srfA* operon, which is unable to produce surfactin (PS-216 Δ*srfA mutant*). Both mutants are reported to still form biofilms in liquid media.^34^ The biofilm formed by both mutants on potato root was barely detectable at 26 hours after inoculation, while strong biofilm formation was observed for wild-type (WT) bacteria (Figure 3A, Figure S6, Table S4). The reduced ability to form biofilm was also reflected in the potato response to *B. subtilis*. The induction of JA synthesis and ethylene signalling markers observed in response to the WT was attenuated in the plant response to the mutants. Similarly, the colonization associated downregulation of apoplastic ROS generator RBOHD and photosynthesis marker CAB occurred to a lesser extent in response to the mutants (Figure 3B, Table S5).

**Figure 3:**
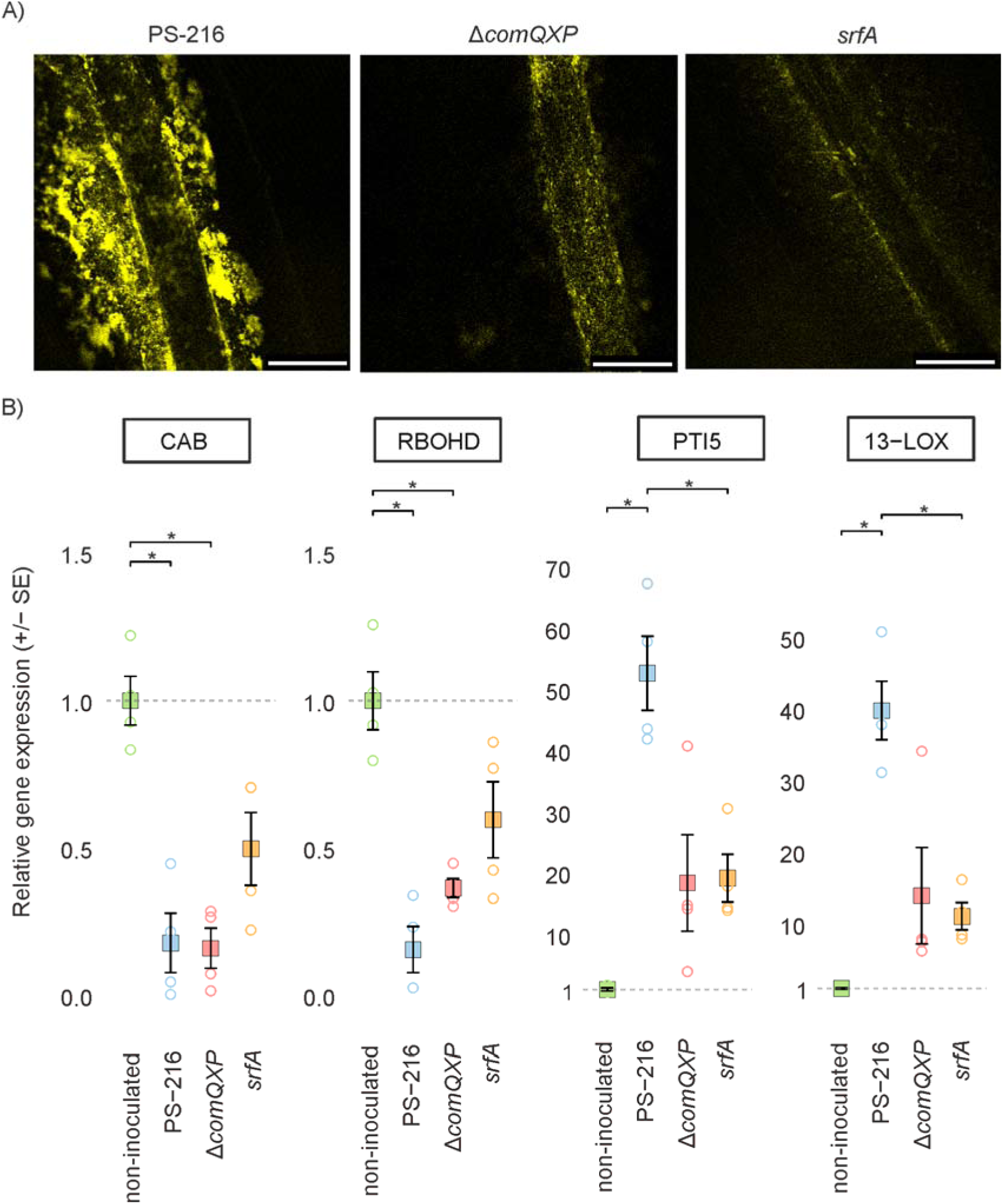
Induction of plant responses is dependent on efficient *B. subtilis* biofilm formation on roots. A) Biofilm formation on potato roots after 26-h incubation with YFP-labelled *B. subtilis* PS-216 (left), quorum sensing-negative *B. subtilis* YFP-labelled mutant PS-216 Δ*comQXP* (middle) and PS-216 srfA mutant (right) culture. Scale is 500 µm. B) Expression of selected genes (RBOHD, 13-LOX, CAB, PTI5) in non-inoculated potato roots and after 26-h incubation with *B. subtilis* PS-216 YFP, PS-216 Δ*comQXP* YFP and PS-216 *srfA* YFP. Relative gene expression was scaled to the average gene expression of non-inoculated group. Welch’s t-test was used to determine differences between groups. Individual measurements (circles), mean (squares) and standard error are shown. Asterisks (*) denote statistically significant difference (p-value < 0.05). See Table S5 for the results of all genes for both roots and shoots.

To confirm that biofilm establishment is required for successful activation of plant responses, and that *B. subtilis* secondary metabolites are not sufficient, we treated plants with the conditioned media in which bacteria were growing. Results showed that the conditioned media alone did not trigger any response in treated plants, albeit containing surfactin (Figure S7, Table S6). Therefore, we conclude that *B. subtilis* biofilm formation on the roots is required for induction of potato response.

### Non-kin intraspecies interaction of *B. subtilis* strains on root intensifies potato responses

To understand plant responses to colonizing bacterial communities, which is relevant for application in practice, we set up an experiment inoculating plants with two genetically different, categorized previously as non-kin, and two isogenic (only differently fluorescently labelled) *B. subtilis* strains. When roots were inoculated with PS-216 and PS-218 strains separately, both equally colonized the roots (Figure 4A, first two columns). Similarly, when differentially labelled isogenic strains were combined, the resulting biofilm on the root contained similar abundances of each strain, with cells of each strain spatially well mixed (Figure 4A, third column). On the other hand, when non-kin strains PS-216 and PS-218 were combined, we observed biofilm with segregation of the two strains (Figure 4A, last column). Moreover, one of the strains dominated on the roots, with only a small amount of biofilm formed by the other strain (Figure 4A, last column).

**Figure 4:**
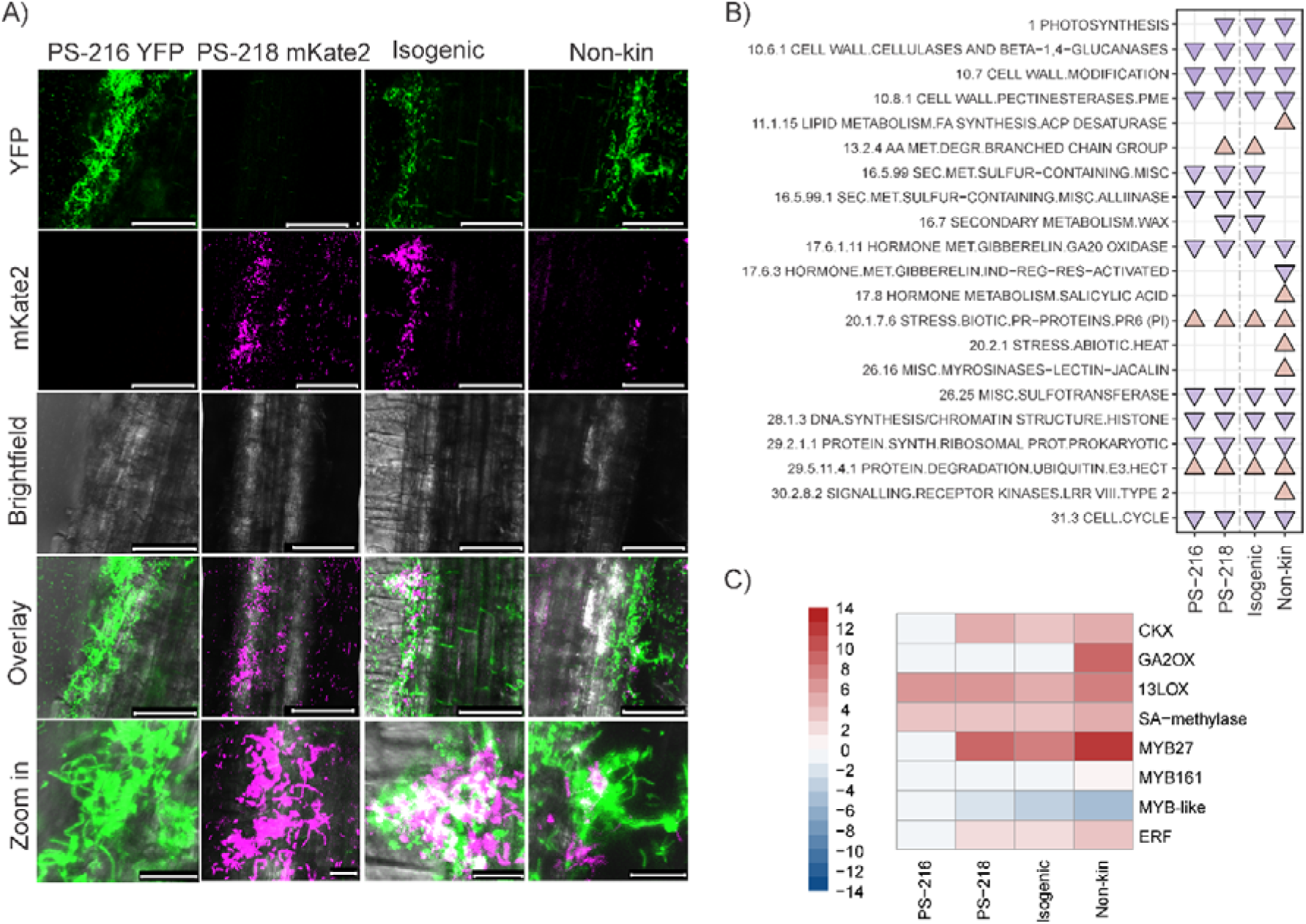
Social interactions of B. subtillis strains on potato roots intensify potato immune response. A) Colonization of potato roots by isogenic and non-kin *B. subtilis* strains after 4-h incubation. From top to bottom: YFP fluorescence, mKate2 fluorescence, brightfield, overlay of images, zoom in of overlay images. From left to right: PS-216 YFP, PS-218 mKate2, PS-218 YFP and PS-218 mKate2 (isogenic – self interaction), PS-216 YFP and PS-218 mKate2 (non-kin interaction). White color (overlay) shows colocalization of YFP and mKate2. The scale is 25 µm for **zoom in** images and 100 µm for all others. B) Visualization of responses in the context of biological pathways. For each comparison between *B. subtilis*-inoculated and non-inoculated plants, gene sets enriched for up- or downregulated genes (q-value < 0.1) are shown and designated as red triangles or blue triangles, respectively after 2-h incubation. MapMan ontology BINs were used to generate gene sets. Sample group comparisons to non-inoculated ones (in columns) are designated with bacterial strain names used for inoculation. PS-216: PS-216-YFP, PS-218: PS-218-mKate2, isogenic interaction: YFP-labelled PS-218 and mKate2-labelled PS-218, non-kin interaction: YFP-labelled PS-216 and mKate2-labelled PS-218. See Table S2 for all results. C) Examples of genes with intensified response when plant is interaction with two non-kin strains. Log2 values are shown. Blue: downregulated genes, red: upregulated genes, according to scale on the left.

We next performed RNA-seq analysis of roots at the time of uniform establishment of *B. subtilis* biofilm (2 hpi). In roots, the intensity of the transcriptional response to the non-kin strain mixture was higher than that to the mixture of two isogenic, differentially labeled strains (Figure 4, Figure S3). Regarding biological pathways, some receptor kinases (BINs 26.16 and 30.2.8.2) were specifically upregulated in the non-kin interaction, as well as some enzymes from fatty acid biosynthesis (BIN 11.1.15) and heat shock proteins (BIN 20.2.1) (Figure 4B, Table S2). In addition, 422 genes had more than 2-fold stronger response in the interaction with non-kin strains compared to monoculture or the isogenic mixture. Among the more induced ones, are two MYB transcription factors, salicylate carboxymethyltransferase, gibberellin 2-oxidase 2, peroxidase and beta-galactosidase genes (Table S3).

### B. subtilis abundance is increased in plants with silenced PTI5

We next overlayed data with plant stress signaling prior knowledge network, built from experimental data on protein–protein interactions, protein–DNA interactions and metabolic pathways (skm.nib.si).^35^ The central signaling module that was regulated in the studied interaction was ethylene signaling, linked to regulated JA synthesis, SA signaling and ROS signaling (Figure 5A; Figure S8; Table S7). Thus, we focused on ERF transcription factors, as the role of ethylene signaling was so far not in depth addressed in interaction of plants with beneficial microbes. Among ERFs, PTI5 was found to be more than 50-fold induced when plants were exposed to *B. subtilis* monocultures or two-strain mixtures, already early after colonization (after 2-hour incubation). This could indicate its role as regulatory hub in plant colonization and/or biofilm formation. To test this role, we produced transgenic potato with reduced expression of PTI5. First, we followed biofilm formation in transgenic plants with silenced PTI5. Results showed that PTI5 is not involved in biofilm formation on roots (Figure 5B, Table S8), as no difference in biofilm formation was detected between non-transgenic (NT) and PTI5-silenced plants (Figure 5B, Figure S9, Table S8).

**Figure 5:**
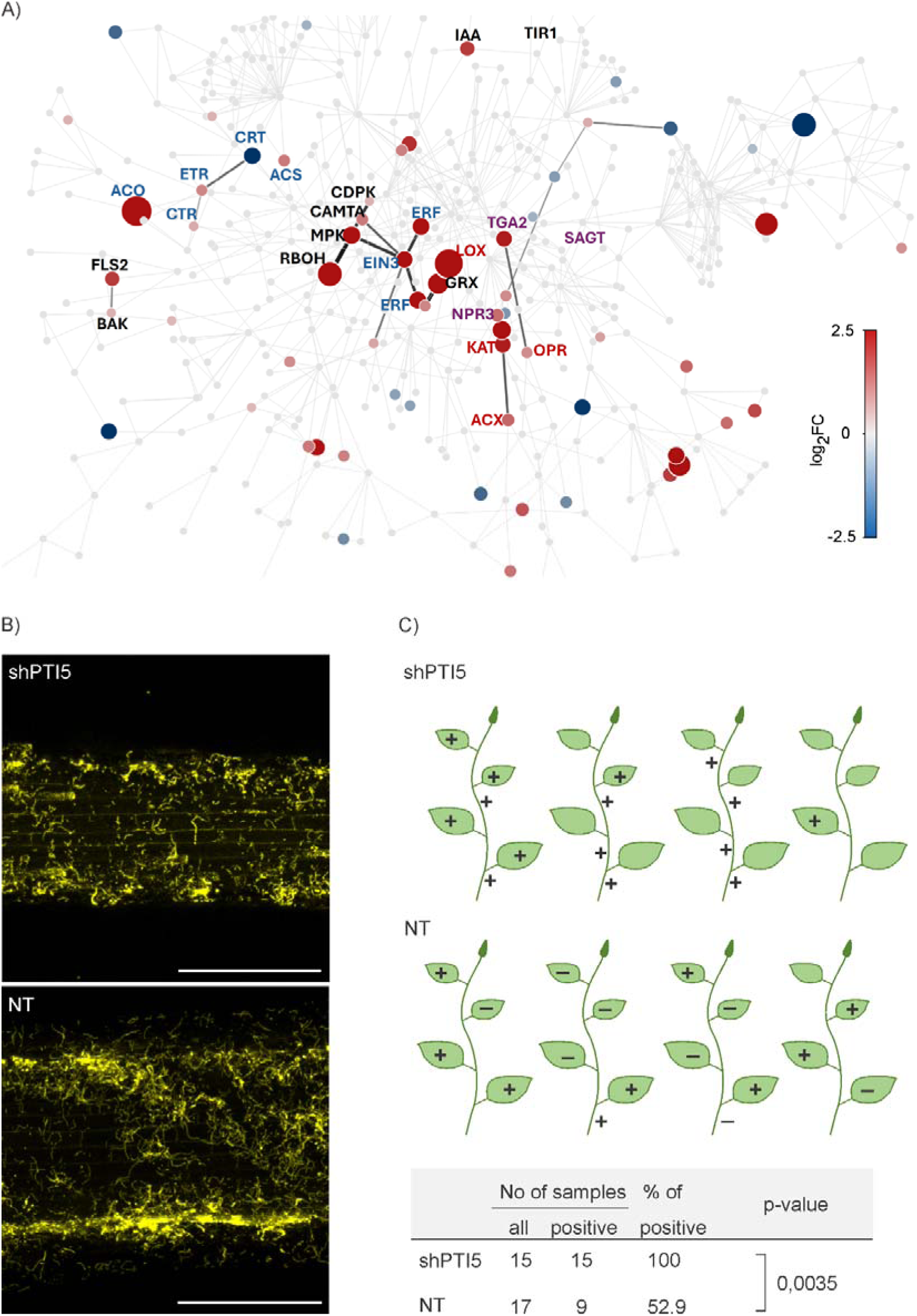
ERF transcription factor PTI5 is required for balanced colonization of potato plants. A) Plant immune signalling modules that were regulated following incubation of roots with non-kin mixture of *B. subtilis* strains. Genes and proteins are represented as nodes and interaction between them (protein–protein interaction, protein–DNA interaction) as edges of the network. Differentially expressed genes from ethylene (blue), JA (red), SA (violoet) and other (black) signalling modules are shown. Black edge connected two differentially expressed genes. All non-differentially expressed genes and edges are shown in grey. The size of the node indicates the difference in expression between PS-216 and PS-218 inoculated and non-inoculated roots (log2FC). See Table S7 for the name and description of regulated genes, and Figure S8 for information on central signalling modules that were regulated in potato roots in isogenic interaction (PS-218 YFP + PS-218 mKate2). B) *B. subtilis* PS-218 YFP root colonization (yellow fluorescence) of transgenic potato with silenced PTI5 (shPTI5) and non-transgenic potato plant (NT) after 2-h incubation. Scale is 250 µm. C) Bacillus abundance in shPTI5 and NT potato plants. The scheme shows leaves of four shPTI5 and four NT plants in which the presence of *B. subtilis* was (+) or was not (-) detected by qPCR in plants from soil which were incubated for 2 h in PS-218 YFP before being planted and grown in soil for one week. Leaves with no mark were not analyzed. The table presents a statistically significant difference in the percentage of positive samples (leaves, nodes and internodes) between NT and shPTI5 plants determined by Fisher’s exact test. See Table S9 for details. The results of line L6 are shown and were confirmed on another transgenic line shPTI5 L2 (Figure S9, Table S10).

PTI5 was also upregulated in potato shoots. To test if it has a role in maintenance of *B. subtilis* endophytic community we followed *B. subtilis* abundance in shoots of NT and PTI5-silenced plants. *B. subtilis* abundance in systemic tissue was higher in PTI5-silenced plants, revealing that the balance in the abundance of endophytic *B. subtilis* community was perturbed (Figure 5C, Table S9, Table S10).

### PTI5 is also regulating the extent of potato root colonization by arbuscular mycorrhizal fungi

Considering the role of PTI5 in regulating endophytic colonization of *B. subtilis*, we aimed to explore if this role was specific for this interaction or could be extended to other beneficial symbiosis. We decided to explore the potential role of PTI5 in regulating the extension of root colonization by arbuscular mycorrhizal fungi. This symbiosis is known to be under plant control to keep the interaction under mutualistic levels and avoid excessive colonization.^36,37^ With this aim, we explored the colonization levels and symbiosis functionality in NT and PTI5-silenced lines. As shown in Figure 6, *R. irregularis* successfully colonized potato roots, with intercellular hyphae, arbuscules and vesicles present in the cortex. Remarkably, the extension of colonization, including the abundance of arbuscules and vesicles was higher in the PTI5-silenced lines (Figure 6A). Indeed, the percentage of root length colonized by the fungus was more than 2-fold higher in the silenced lines (Figure 6B), in agreement with the higher amount of fungal DNA in those plants. To check if the functionality of the mycorrhizal symbiosis was altered in those lines, we checked the expression levels of plant and fungal transporters involved in the nutrient exchange between the symbionts within the arbuscules: the potato phosphate transporter StPT4, and the fungal monosacharide transporter RiMST2, both transcriptionally upregulated in arbusculated cells and defined as markers for symbiotic functionality (Figure 6C). Both markers were upregulated in the PTI5-silenced lines, confirming the functionality of the symbiosis.

**Figure 6:**
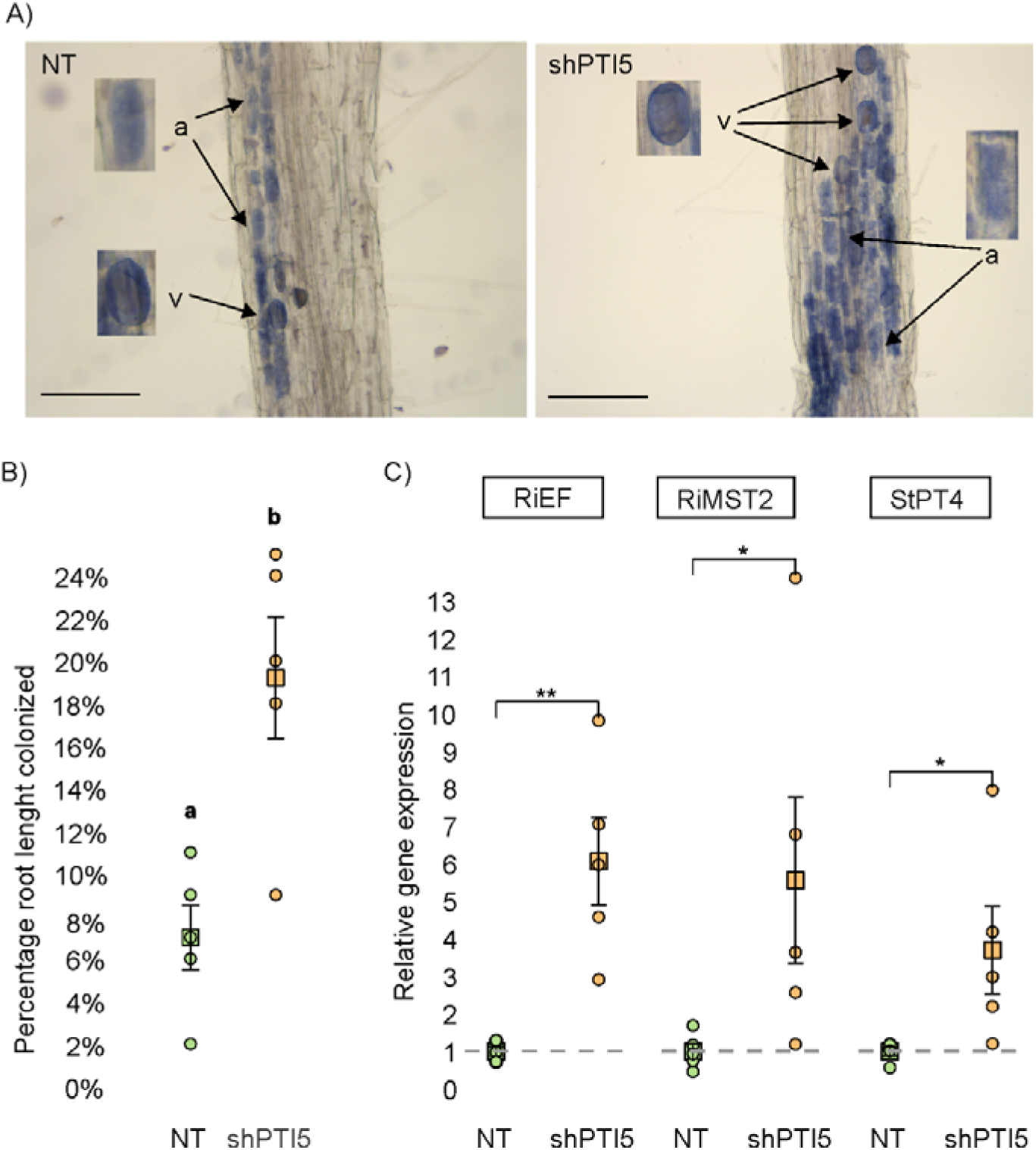
Mycorrhizal colonization of potato roots in non-transgenic potato plant (NT) and PTI5-silenced genotype (shPTI5). A) Representative images of mycorrhizal colonization by *R. irregularis* in the different genotypes. Fungal structures within the root cortex of potato plants are stained in blue (a: arbuscules, v: vesicles). Scale is 200 µm. B) Percentage of root length colonized by the arbuscular mycorrhizal fungi *R. irregularis*. C) Expression analysis of *R. irregularis* constitutive gene (RiEF) and symbiosis marker genes, the *R. irregularis* monosaccharide transporter (RiMST2) and the potato phosphate transporter (StPT4), as well as regression analysis using Dunn’s pairwise comparisons were used to determine the differences between the genotypes. Different letters show statistical differences. Permutation test was used to determine the differences between the genotypes. Individual measurements (circles), mean (squares) and standard error are shown. Asterisks denote a statistically significant difference (*: p-value < 0.05, **: p-value < 0.01). The results of analysis of line L6 are shown and were confirmed on another transgenic line shPTI5 L2 (Figure S10).

## DISCUSSION

Microbial inoculants have been proven as efficient and environmentally friendly alternatives to chemical pesticides and fertilizers.^38,39^ Current limitation in adopting such agricultural practices is their variable efficiency in the field due to lack of understanding of mechanisms involved in the regulation of plant colonization and maintenance of the plant microbiome.^6,40^ Therefore, studying the mechanisms regulating the interactions between crop plants and microbes is crucial to ensure robustness of such plant protection approach. As the third most important crop in the world, exceptionally sensitive to a wide range of stresses, potato is one of the most suitable candidates for plant growth-promoting rhizobia (PGPR) based products.^31^ We here established a model system for such studies, exploring selected *B. subtilis* soil isolates individually or in mixtures. Furthermore, we explored the potential extension of our findings by testing other widespread mutualistic interaction, the arbuscular mycorrhizal symbiosis.

*B. subtilis* initially suppresses immune response in potato roots (e.g. downregulation of diverse receptor kinases) thereby facilitating its own colonization, while it boosts induction of plant immunity related genes in shoots at a later stage (Figure 2, Table S2), as it was shown also in other studies of plants with beneficials as part of induced resistance phenomena.^13,41^ Here we show that live bacteria are required for triggering of the plant response, as the conditioned media in which bacteria were growing did not trigger the response (Figure S7). Moreover, the intensity of the plant response was dependent on the extent of biofilm formation on the potato root (Figure 3). Both, the mutant in surfactin production and mutant in QS (also deficient in surfactin production) showed a reduced ability to form biofilm on roots. This is interesting as the same strain was able to form floating biofilms (pellicles),^42^ suggesting that the ComX QS control of the biofilm development depends also on plant factors. Besides residing on the plant roots, *B. subtilis* can also colonize the plant endophytically. Internalised bacteria stop dividing and change the morphology (Figure 2, Figure S5). The round bacterial form closely resembles L-form bacteria, which are characterized by modified or absent cell walls.^42,43,44^ We also tested the extent of bacteria internalisation with QS impaired strains, which was interestingly not affected (Table S11).

To get insights into the complexity of plant–microbe interactions, we extended our studies to application of two different *B. subtilis* strains to explore how bacterial diversity may influence the plant response. We studied pairs of strains with different kin discrimination types of intraspecies interactions — isogenic strains were predicted to be cooperative, and non-kin strains to be antagonistic.^22^ We observed that non-kin interaction of two *B. subtilis* strains invigorates the plant response, especially in immune and abiotic signalling components (Figure 4, Table S3). These results imply that even minor taxonomic changes in the phytobiome can lead to substantial differences in the plant responses. Increasing evidence support that plants actively manage their microbiota for balancing growth with immunity.^46^ It has been reported that *B. subtilis* can promote plant growth via direct pathways such as producing phytohormones (e.g., auxin, gibberellin, cytokinin, and abscisic acid).^46,47,48^ Both strains used in this study produce auxins (Table S12). Interestingly, we here show that *B. subtilis* additionally regulates phytohormone metabolism *in planta*. Gibberellin 2-oxidase 2 and cytokinin oxidase, which catalyze the irreversible degradation of cytokinin and gibberellin phytohormones, have been among the most induced genes by inoculation of non-kin *B. subtilis* strains (Figure 4, Table S3). This could suggest that equilibrium between growth and immunity is shifted towards immunity in plants inoculated by non-kin *B. subtilis* strains, as a functional relation between lower activity of gibberellin signalling and reduced plant disease severity has previously been confirmed in potato.^50^ In accordance, indole-3-acetic acid-amido synthetase GH3.8 was in our study induced in *B. subtilis* inoculated plants at late time point in both shoots and roots (Table S3). GH3.8 functions as IAA-amino synthetase that prevents the accumulation of free indole acetic acid (IAA), which increases plant disease resistance, growth and development.^51^

Not much is however known on the mechanisms that allow for successful colonisation of plants with beneficial microbes while avoiding colonization by harmful pathogens. MTI is known to be induced also when plant is exposed to beneficials.^11^ In interaction of Arabidopsis with beneficial *Pseudomonas* strain it was found that suberin biosynthesis is down regulated to allow for better penetration of bacteria.^11^ Similarly, in our study cell wall metabolism was downregulated in root colonization (Figure 2, Figure 4). In addition, synthesis of toxic terpenoids was downregulated in roots at the biofilm forming stage (Figure 2). Suppression of toxic ROS is crucial for successful colonisation of plants by microbes.^24^ We found ROS scavenger peroxidase and ROS generator RBOHD induced and repressed, respectively, following inoculation by *B. subtilis* (Figure 3). Several studies also reported the increase of peroxidase in different plant–*Bacillus* interactions, and in some cases this induction have been associated to pathogen resistance.^51,52,53,54,55,56^

Plant colonisation is frequently linked to induction of SA, JA and ethylene signalling pathways and transcriptional rewiring, of which MYB72, MYC2 and NPR1 transcription factors are the most studied ones, and are proposed to be relevant for IR responses.^13,58^ Here we found upregulation of the 13-LOX pathway leading to JA synthesis, as previously described in other plant-PGPR interactions (Figure 1, Figure 3).^58,59,60^ Similarly, several studies reported activation of SA signalling pathway.^14^ We found NPR1 upregulated in shoots (Table S2, Table S3). In corroboration with previous studies, we also identified several MYB transcription factors induced in both roots and shoots (Table S3).

Modulation of ethylene signalling in plant microbiome interactions have been pinpointed as a central element in their regulation and their impact on plant responses to stress.^62^ For example, ethylene have been proposed to regulate biofilm formation and root colonization by beneficial bacteria^63,64^ and the arbuscular mycorrhizal symbiosis,^65^ and it is required for IR by beneficial bacteria and fungi.^66,67^ We also detected induction of the ethylene signalling pathway in *B. subtilis* inoculated roots, including ETR, EIN3 and several ERF transcription factors (Figure 5, Figure S8, Table S3). Moreover, 1-aminocyclopropane-1-carboxylate oxidase, which catalyzes the final step in ethylene biosynthesis, was in our study induced in all tissue and time points, further suggesting involvement of ethylene signalling pathway in potato response to *B. subtilis* (Figure 5, Figure S8, Table S3).

ERF transcription factors, the members of the AP2/ERF family active under ethylene signling, play prominent roles in plant immunity, although the function of most of them remains unknown.^13,59,68,69^ A different set of ERFs was regulated in roots and shoots in our study. Among them, ERF transcription factor PTI5 was strongly upregulated in all studied conditions. PTI5 was first identified in tomato as interacting partner of the *R* gene *Pto* that confers resistance against the bacterial pathogen *Pseudomonas syringae* pv. tomato.^70,71^ On the other hand, we have previously shown that PTI5 negatively regulates potato defence response to pathogens potato virus Y and *Ralstonia Solanacearum*, suggesting its role as susceptibility factor in potato immunity.^33^ Interestingly, PTI5 is transcriptionally regulated by EIN3 in the presence of SA only and on the protein level regulated by autophagy degradation.^33^ Results of this study suggest that PTI5 has a negative regulatory role also in the maintenance of endophytic community in potato, as both *B. subtilis* abundance in systemic tissues and mycorrhizal colonization of roots were increased in PTI5-silenced potato plants (Figure 5, Figure 6). High PTI5 levels thus both increase susceptibility of potato to pathogens and reduce the capability of plant to interact with beneficial organisms. PTI5 thus represents a target for future breeding efforts leading towards sustainable potato production. Taking it altogether, we here set a cornerstone in understanding how beneficial microbes and plant communicate to regulate endophytic colonization and maintain mutualism and have identified a way for improved colonization by beneficial through silencing of PTI5.

## ACKNOWLEDGEMENTS

We thank Nastja Marondini, Anja Moškrič, Tina Koželj, Tim Godec, Ana Žuran and Olena Nesterenko for technical support and laboratory assistance. This research was financially supported by the Slovenian Research Agency grants P4-0165, P4-0116, J4-9302, J4-4550 and J4-3089, and grant MCIN/AEI/ PID2021-124813OBC31, “ERDF A way of making Europe” financed by the European Union.

## AUTHOR CONTRIBUTIONS

Concept and design: TL, MJP, IMM, KG. Acquisition, analysis or interpretation of data: TL, BK, KP, KS, TGK, MZ, MP, PS, AV, VL, TMP, JMP, MJP, IMM, KG. Drafting of manuscript: TL, KG. Critical revision of the manuscript for important intellectual content: MZ, MP, PS, MJP, IMM, KG.

## METHODS

### Plant material

Potato plants cv. Rywal (NT) and shPTI5-Rywal were grown in stem node tissue culture for two weeks after *in-vitro* micropropagation under controlled environmental conditions (22⍰±⍰2°C in the light and 19⍰±⍰2°C in the dark with 70–90 μmol/m2/s2 radiation (OSRAM L 58 W/77 FLUORA lamps, Germany) and a 16-h photoperiod) as previously explained.^72^ For studying systemic *B. sublilis* spread, plants were planted in soil two weeks after node segmentation and grown under controlled environmental conditions for one week as previously described.^73^ shPTI5-Rywal transgenic lines were prepared by introducing shPTI5 construct^33^ into NT using *A. tumefaciens* LBA4404.^74^ The transformed bacteria were used for the transformation of NT stem internodes from *in⍰vitro* plantlets as described in Lukan et al.^73^ using appropriate selection media.

### B. subtilis strains

WT *B. subtilis* strains PS-216 and PS-218^75^ tagged with red fluorescent protein (mKate2) and yellow fluorescent protein (YFP) gene linked to a constitutive hyperspank promotor were used, namely PS-218 *amyE::P_hypercl03_-YFP* (Sp) (PS-218 YFP), PS-216 *amyE::P_hypercl03_*-*YFP* (Sp) (PS-216 YFP), PS-218 *amyE::P_hyperspank_-mKate2* (Chl) (PS-218 mKate2) and PS-216 *amyE::P_hyperspank_-mKate2* (Chl) (PS-216 mKate2).^20,22^ PS-216 mutants with impaired surfactin production PS-216 *amyE::PhypercI03-YFP* (Sp) *srfA::Tn917 (mls)* (PS-216 *srfA* YFP)^22^ and QS mutant PS-216 *ΔcomQXP::kan amyE::P -YFP* (Sp) (PS-216 Δ*comQXP* YFP) (this work) were used. QS mutant PS-216 ΔcomQXP::kan amyE::PhypercI03-YFP (Sp) was constructed by transformation of PS-216 amyE::PhypercI03-YFP (Sp)^22^ with plasmid pED302,^76^ carrying a kanamycin cassette between *degQ* and *comA*, resulting in a deletion of the *comQXP* locus.

### IAA production detection

The production of IAA was assessed following the protocol of Pramanik et al.^77^ Fresh bacterial cultures were prepared from single colonies in LB medium supplemented with 0.1 % tryptophan and incubated in the dark at 30°C with shaking at 120 rotations per minute (rpm) for 48 hours. After incubation, cultures were centrifuged at 10,000 rpm for 5 minutes, and the supernatant was transferred to a microtiter plate in triplicates. Equal volumes of Salkowski reagent (prepared by adding 1 mL of 0.5 M ferric chloride to 50 mL of 35 % perchloric acid, mixed well, and stored in a brown bottle) were added to each well. The mixture was incubated in the dark at room temperature for 30 minutes. Blanks were prepared similarly, using only LB medium with 0,1 % tryptophan. Absorbance was measured spectrophotometrically at 536 nm, and IAA concentrations were calculated using a standard curve prepared with concentrations ranging from 10–100 µg/mL of IAA.

### Potato inoculation with *B. subtilis*

On day one, *B. subtilis* from permanent culture was plated on a solid LB medium with appropriate (see methods section *B. subtilis* strains) antibiotics (100 µg/mL spectinomycin or 5 µg/mL chloramphenicol) and incubated overnight at 37°C. On day two, a single colony was resuspended in 3 mL of LB medium with antibiotics and grown overnight at 200 rpm and 37°C. On day three at noon, 30 µL of the bacteria culture was inoculated in 3 mL of LB medium without antibiotic and grown for 3 hours at 200 rpm at 37°C (OD_600_ between 0.4 and 0.6). Bacterial culture was then diluted in 3 mL of MS30 (5 g of Murashige and Skoog Basal Medium with vitamins and 30 g of sucrose per 1 L of bidistilled water, pH 5.8) medium to OD_600_ 0.02.

For studying marker genes expression after potato incubation in bacterial culture of low density, two weeks old potato plants from tissue cultures were added to diluted bacterial culture. Note that diluted bacterial culture was incubated 16 hours at 11 rpm before adding potato plants. Shoots were separated from the culture by parafilm cover and incubated for 21 hours at 11 rpm in a growth chamber prior sampling for quantitative PCR (qPCR) (Exp1 and Exp2 in Figure S2). When studying if the conditioned media triggered any response in treated plants (Table S6), plants were added to a culture (shoots were separated from the culture by parafilm cover) and discarded after 17-hour incubation. New plants were added to conditioned medium and incubated for 26 hours.

For studying *B. subtilis* morphology after internalization in plants, two weeks old potato plants in tissue cultures grown in the solid MS30 medium were inoculated with 300 µL of bacterial culture, which was spread on the medium and after a specified time (Figure S4), stem and leaves were observed under confocal microscope.

In all other experiments, on day four in the morning (after 16-hour incubation), bacterial cultures were vortexed, except when studying the role of QS and surfactin (Table S5, Table S6). Cultures of different strains were mixed in a ratio of 1:1 when interactions were studied. In 3 mL of bacterial cultures, two-week old potato plants from tissue cultures were added (shoots were separated from the culture by parafilm cover) and incubated for a specified time (15 min to 26 hours), depending on the experiment. When studying *B. subtilis* abundance in shPTI5 transgenic plants with qPCR (Table S9, Table S10), plants were planted in soil after a 2-hour incubation. In all experiments, shoots were not in contact with bacterial culture.

### Mycorrhizal symbiosis establishment and quantification

The mycorrhizal inoculum was maintained as monoxenic culture consisting in an *in vitro* culture of transformed carrot roots colonized by the arbuscular mycorrhizal fungus *R. irregularis* (MUCL 57021) (REKA Group. B.V., Bleiswijk, The Netherlands), produced and maintained growing in Gel-Gro (ICN Biochemicals, Aurora, OH, USA) as previously described in Chabot et al.^78^ Potato plantlets growing in 125 mL pots with a mixture of soil, sand and vermiculite (3:2:1, v:v:v) were inoculated by adding a 1 cm piece of the monoxenic culture, containing approximately 50 *R. irregularis* spores, fungal hyphae and colonized carrot roots. In the non-inoculated treatment, a piece of medium containing only uninfected carrot roots was applied to the non-mycorrhizal controls. Plants were grown in a growth chamber (day/night cycle: 16 h, 24°C/8 h, 19°C; relative humidity: 50 %). The experiment was repeated twice and eight independent plants per genotype were analyzed.

Mycorrhizal colonization of roots and symbiosis establishment was evaluated histochemically and molecularly mycorrhizal structures within the roots were observed and quantified after clearing the roots in 10 % KOH and staining the fungal structures with 5 % black ink in 2 % acetic acid solution as described in García at al.^79^ Percentage of mycorrhizal colonization of roots was determined following the gridline intersection method^80^ using a Nikon SMZ1000 stereomicroscope. To quantify the total amount of fungus within the root and assess symbiotic functionality we analysed the expression levels of marker genes by qPCR (see below).

### Confocal microscopy

Stellaris 8 Confocal microscope with HC PL FLUOTAR 10x/0.30 DRY objective (Leica Microsystems) was used to detect emission of YFP and mKate2 on the roots of plants in the experiment for investigating timing of biofilm formation, *B. subtilis* morphology after internalization in plants, biofilm formation pattern when studying bacteria–plant social interactions, biofilm presence on the roots of plants sampled for RNA-seq, biofilm formation in transgenic plants with silenced PTI5 and prior all qPCR experiments to confirm roots colonization. Confocal microscope Leica TCS LSI macroscope with Plan APO 5 x objective (Leica Microsystems) was used to detect emission of YFP when investigating biofilm formation on the roots of mutants that have attenuated surfactin production. The emission of YFP was followed after excitation with 488 nm laser in the window 505–560 nm. The emission of mKate was followed after excitation with 589 nm laser in the window between 605–650 nm. Regions of interest (ROI) were scanned unidirectionally with scan speed of 400 Hz and frame average at least 2. The images were processed to obtain maximum projections from Z-stacks for all channels and merged using Leica LAS X software (Leica Microsystems). Mean intensity of fluorescence on confocal images was determined in LAS X software (Leica). Statistically significant differences were determined by Student’s t-test.

### Transmission electron microscopy (TEM)

Samples were prepared for examination with TEM using negative staining method. Pellets of centrifuged liquid culture or leaf homogenate infiltrated with *B. subtilis* were resuspended in 0.1 M phosphate buffer with 2 % polyvinylpyrrolidone. Six µL of resuspension was applied to copper grids with or without glow discharge (400 mesh, formvar-carbon coated) for 5 minutes. Afterwards, the remaining resuspension was dabbed with filter paper. This procedure was repeated two more times.

Afterwards, the grids were washed with bidistilled water and stained with one droplet of 1 % (w/v) water solution of uranyl acetate. Two grids were prepared for each sample. The grids were observed by transmission electron microscope TALOS L120 (ThermoFisher SCIENTIFIC, The Netherlands), operating at 100 kV and representative micrographs were acquired (camera Ceta 16M) using Velox software.

### qPCR analysis

The shoots and elongation zones of roots of *B. subtilis*-inoculated and non-inoculated (see description of inoculation chapter) potato plants were sampled in four or five biological replicates (plants). RNA was isolated by RNeasy Plant Mini Kit (Qiagen) from shoots and RNeasy Plant Mini Kit (Qiagen) or RNeasy Micro Kit (Qiagen) from roots according to the manufacturer’s protocols. Concentration and purity of DNase-treated (0.51μL of DNase I per μg RNA; Qiagen) total RNA was evaluated using NanoDrop ND1000 spectrophotometer (Nanodrop technologies) and agarose gel electrophoresis and then reverse-transcribed using the High-Capacity cDNA Reverse Transcription Kit (Thermo Fisher Scientific) or High-Capacity RNA-to-cDNA Kit (Thermo Fisher Scientific). To determine *B. subtilis* abundance in systemic shoot tissue, genomic DNA was isolated by MagMAX™ Plant DNA Isolation Kit (Thermo Fisher Scientific) according to the manufacturer’s protocol. Concentration and purity were evaluated as stated above.

Samples were analyzed in the setup for qPCR as previously described.^32^ The expression of 8 genes (RBOHD, HSP70, PTI5, 13-LOX, PR1B, BGLU2, CPI8, CAB) involved in different steps of immune signaling (Table S13 for a full list of genes and their description) was determined and normalized to the expression of two validated reference genes, COX and elongation factor 1 (EF-1), as described previously.^81^ See Lukan et1al.^32^ for primer and probe information. Relative copy numbers of *comQ* from *B. subtilis* were determined and normalized to relative copy numbers of NPR1 (a single copy gene, https://spuddb.uga.edu/). The standard curve method was used for relative gene expression quantification using quantGenius (http://quantgenius.nib.si).^82^

For molecular assessment of mycorrhizal colonization, the expression of the constitutive *R. irregularis* elongation factor 1 (RiEF) gene and the markers of symbiotic functionality, the fungal RiMST2 and the plant StPT4, both involved in the nutrient exchange between the symbionts in the arbuscules were determined in roots by qPCR using gene-specific primers (Table S13). Relative quantification of specific mRNA levels was performed using the comparative 2-Δ(ΔCt) method.^83^ Expression values were normalized using the normalizer gene EF-1α encoding the potato translation elongation factor-1α. ^84^ Five independent biological replicates per treatment were analyzed.

Statistical analysis was conducted in R version 4.3.1.^85^ Relative gene expression was scaled to the average relative gene expression of the non-inoculated group.^86^ Mean, standard deviation, standard error of the mean and 95 % confidence intervals were calculated per gene, tissue type and genotype and visualised using the ggplot2 package v3.3.6.^87^ Pairwise Welch’s t-test with holm correction for multiple testing adjustment was employed to quantify the difference between the groups. Results were visualised using rstatix v0.7.0^88^ and ggpubr v0.4.0^89^ R packages.

### RNA-seq analysis

RNA was isolated from shoots and elongation zones of roots by RNeasy Plant Mini Kit (Qiagen) and treated with RNA Clean & Concentrator-5 (DNase Included) (Zymo Research). In RNA-seq Exp 1, RNA-seq was for roots performed after 2- and 26-hours incubations with *B. subtilis* culture and for shoots after 26 hours of incubation, both for monocultures (PS-216 YFP, PS-218 YFP) and non-inoculated samples (incubation in MS30 medium) as a control. In RNA-seq Exp 2, RNA-seq was for roots performed after 2 hours of incubation with *B. subtilis* culture for monocultures (PS-216 YFP, PS-218 YFP, PS-218 mKate2), isogenic mixture (PS-218 mKate2 + PS-218 YFP), non-kin mixture (PS-218 mKate2 + PS-216 YFP) and non-inoculated samples (incubation in MS30 medium) as a control. Four biological replicates (plants) were analyzed for each interaction (Table S14).

Raw paired-end Illumina reads were adapter-trimmed and reads below average PHRED basecall quality score 20 were removed using Trim Galore (https://github.com/FelixKrueger/TrimGalore).^90^ Sequencing quality control was performed using FastQC v0.11.9.^91^ Taxonomic classification of reads was performed using Centrifuge v1.0.4^92^ in conjunctions with the 2018 nt database. Reads were mapped to *Solanum tuberosum* Phureja clone DM 1-3 genome with STAR v2.7.5c^93^ using the merged genome annotations GFF3.^94^ Only uniquely mapped fragments were counted (-- outFilterMultimapNmax 1) and additional alignment parameters were set to restrict erroneous mapping (--outFilterMismatchNoverReadLmax 0.02, --quantTranscriptomeBan Singleend, -- outFilterType BySJout, --alignSJoverhangMin 10, --alignSJDBoverhangMin 1, --alignIntronMin 20, -- alignIntronMax 10000, --alignMatesGapMax 10000). Centrifuge and mapping results showed substantial contamination of sample BS472, therefore we excluded this sample from further analysis.

Gene Set Enrichment Analysis^95^ was performed using MapMan^96^ based gene sets (gmt file available at https://github.com/nib-si/ptda)^97^ and non-filtered normalized read counts, comparing different Bacillus treatments to non-inoculated plants to identify significantly altered regulation of processes and functionally related gene groups (FDR corrected q-value < 0.10). Results of all GSEA comparisons were merged using R v4.3.1.^85^ GSEA figure was generated using adjusted gseaFromStats function from biokit v0.1.1 package.^98^

Differential gene expression analysis was performed in R using the *limma* package.^99^ Low expressed genes were filtered by retaining only genes with raw read counts above 100 in at least 11 samples for RNA-seq Exp 2 and above 50 counts in at least 4 samples for RNA-seq Exp 1. Root and leaf samples were analyzed separately. Normalized counts were transformed using voom function, followed by linear model fit, and statistics calculation for defined contrasts using the eBayes function. Genes with Benjamini-Hochberg FDR adjusted p-values < 0.05 and |log_2_FC| > 1 were considered significantly differentially expressed. Venn diagrams were plotted using a modified limma::vennDiagram function. Raw and normalized read counts were deposited in GEO under accession number GSE232028. Results of RNA-seq were validated by qPCR (Table S1).

### Network analysis

Mechanistic Plant Stress Signalling (PSS) (https://skm.nib.si/)^35^ reaction network was pre-processed using DiNARs’^100^ pre-processing app. Interactions among different molecular entities (such as genes, proteins or transcripts) are defined on functional cluster level, mostly encompassed of *A. thaliana* gene identifiers. Therefore, *Solanum tuberosum* gene identifiers^94^ were first translated to Arabidopsis gene identifiers using BLAST+ reciprocal best hit (RBH) search^101^ on proteomes (file available at https://github.com/NIB-SI/DiNAR/tree/master/TranslationTables). Top three potato identifiers per Arabidopsis identifier were considered and further prioritised to the functinal cluster using DiNARs’ prioritisation script. Experimental data were superimposed onto the PSS network and visualised using DiNAR. Separate condition specific networks, as defined by differential expression contrasts, were exported as individual interactive .html files for further examination.

## Supplemental information

Figure S1: Biofilm on potato roots after incubation in *B. subtilis* culture, imaged at different time points.

Figure S2: Expression profiles of selected genes.

Figure S3: Number of genes, regulated by *B. subtilis* in plants incubated in *B. subtilis* culture of high density (107 CFU/m^L^).

Figure S4: Observation of *B. subtilis* cells in potato tissue after internalization.

Figure S5: B. subtilis cells as observed in liquid culture (A) and potato leaf homogenate (B).

Figure S6: Biofilm formation on potato roots inoculated with *B. subtilis* mutants that have attenuated surfactin production.

Figure S7: *B. subtilis*-produced secondary metabolites in the medium do not contribute significantly to the PTI5 and 13-LOX upregulation.

Figure S8: Regulated central signaling modules in potato roots in isogenic and non-kin interactions.

Figure S9: *B. subtilis* root colonization of transgenic potato lines with silenced PTI5 (shPTI5 L2 and L6) and non-transgenic (NT) potato plant.

Figure S10: Mycorrhizal colonization of potato roots in non-transgenic potato plant (NT) and PTI5 silenced genotype line 2 (shPTI5).

Table S1: Standardised qPCR gene expression data.

Table S2: Results of gene set enrichment analysis.

Table S3: Differential gene expression table for comparisons between plants inoculated with *B. subtilis* and non-inoculated plants.

Table S4: The amount of biofilm on potato roots.

Table S5: Relative gene expression in potato inoculated with *B. subtilis* mutants that have attenuated surfactin production.

Table S6: Expression of selected potato genes induced by conditioned medium in which *B. subtilis* was growing.

Table S7: Plant stress signaling (PSS) prior knowledge network node information with corresponding RNA-seq differential expression.

Table S8: The amount of biofilm formation on potato roots for non-transgenic (NT) and shPTI5 plants.

Table S9: *B. subtilis* abundance in shPTI5 L6 and non-transgenic (NT) potato plants determined by qPCR.

Table S10: *B. subtilis* abundance in shPTI5 L2 and non-transgenic (NT) potato plants determined by qPCR.

Table S11: Abundance of *B. subtilis* PS-216 and QS impaired strain PS-216 *srfA* in Rywal non-transgenic (NT) potato plants, determined by qPCR.

Table S12: The production of indole acetic acid (IAA) was assessed following the protocol of Pramanik et al.^77^

Table S13: Selected genes and corresponding primers and probes sequences for expression analyses with quantitative PCR.

Table S14: Sample information for RNA-seq experiments.

