## Supplemental information for "An ERF transcription factor PTI5, a novel regulator of endophyte community maintenance in potato"

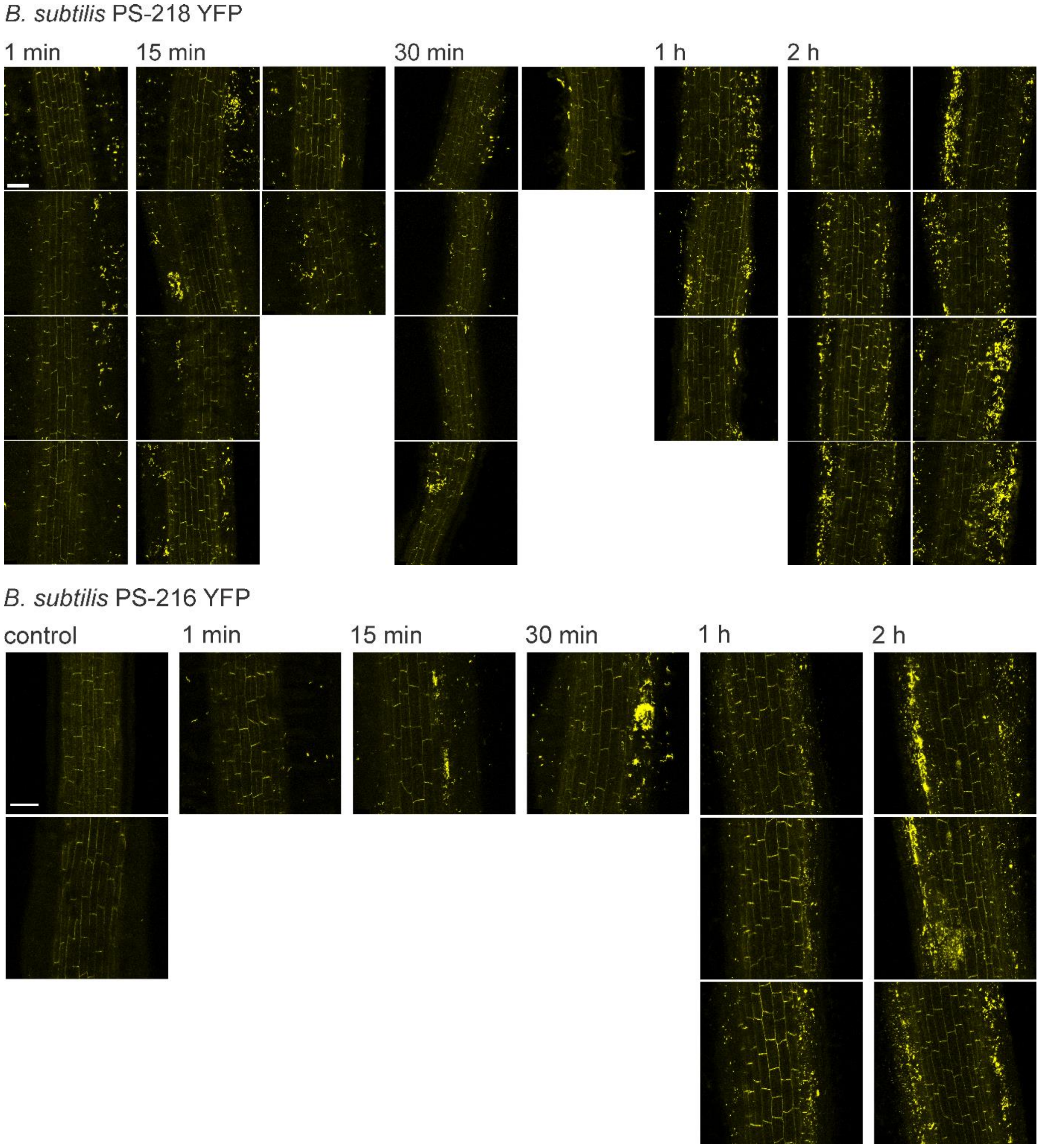


**Figure S1: Biofilm on potato roots after incubation in *B. subtilis* culture, imaged at different time points.** Biofilm formation on potato cv. Rywal roots was imaged by following YFP fluorescence of YFP-tagged *B. subtilis* PS-218 or PS-216 culture after 1 min, 15 min, 30 min, 1 h and 2 h incubation in bacteria culture of high density ($10^7$ CFU/mL). YFP fluorescence is coloured as yellow. Note that cell walls are visible in YFP channel due to autofluorescence. Scale is 100 µm.

| | Test | | Exp1 | | | Exp2 | | | | |
|---|---|---|---|---|---|---|---|---|---|---|
| | PS-216 | PS-218 | PS-218 | | | PS-218 | | | | |
| | | | 2 h | 4 h | 6 h | 1 min | 15 min | 30 min | 1 h | 2 h |
| **Shoots** | | | | | | | | | | |
| RBOHD | ↑ | ↑ | na | na | na | na | na | na | na | na |
| PR1B | ↑ | ↑ | na | na | na | na | na | na | na | na |
| CPI8 | ↑ | ↓ | na | na | na | na | na | na | na | na |
| CAB | ↑ | ↑ | na | na | na | na | na | na | na | na |
| BGLUII | ↑ | **↑** | ↑ | ↑ | ↑ | na | na | na | na | na |
| PTI5 | **↑** | **↑** | ↑ | ↑ | **↑** | na | na | na | na | na |
| HSP70 | ↑ | **↑** | ↑ | ↑ | ↑ | na | na | na | na | na |
| 13LOX | ↑ | ↑ | ↓ | ↑ | ↑ | na | na | na | na | na |
| **Roots** | | | | | | | | | | |
| RBOHD | ↓ | ↑ | na | na | na | na | na | na | na | na |
| PR1B | ↑ | ↑ | na | na | na | na | na | na | na | na |
| CPI8 | ↑ | ↑ | na | na | na | na | na | na | na | na |
| CAB | ↑ | ↑ | na | na | na | na | na | na | na | na |
| BGLUII | ↑ | ↓ | ↑ | ↑ | ↑ | ↑ | ↓ | ↑ | ↑ | ↓ |
| PTI5 | **↑** | ↑ | ↑ | ↑ | **↑** | ↓ | ↑ | ↑ | ↑ | **↑** |
| HSP70 | **↑** | ↑ | ↑ | **↑** | ↑ | ↑ | **↓** | ↑ | ↑ | ↑ |
| 13LOX | **↑** | **↑** | **↑** | **↑** | **↑** | ↓ | ↓ | ↑ | **↑** | **↑** |

**Figure S2: Expression profiles of selected genes.** Expression profiles were determined for shoots and roots of potato plants of cv. Rywal after potato plantlets from tissue culture were added to YFP-tagged *B. subtilis* (strains PS-216 and PS-218) culture with OD600 0.02 and incubated overnight (Test), or were added to *B. subtilis* culture after bacteria have reached $10^7$ CFU/mL and incubated for a specified time (Exp1, Exp2). Expression profiles (up- or downregulation) were determined by comparing expression in inoculated plants with expression in non-inoculated plants (incubated in MS30 medium). ↑: upregulation, ↓: downregulation, **↑**: statistically significant upregulation determined by Welch's t-test, **↓**: statistically significant downregulation determined by Welch's t-test. Exp: experiment, na: not analysed. See Table S1 for all data.

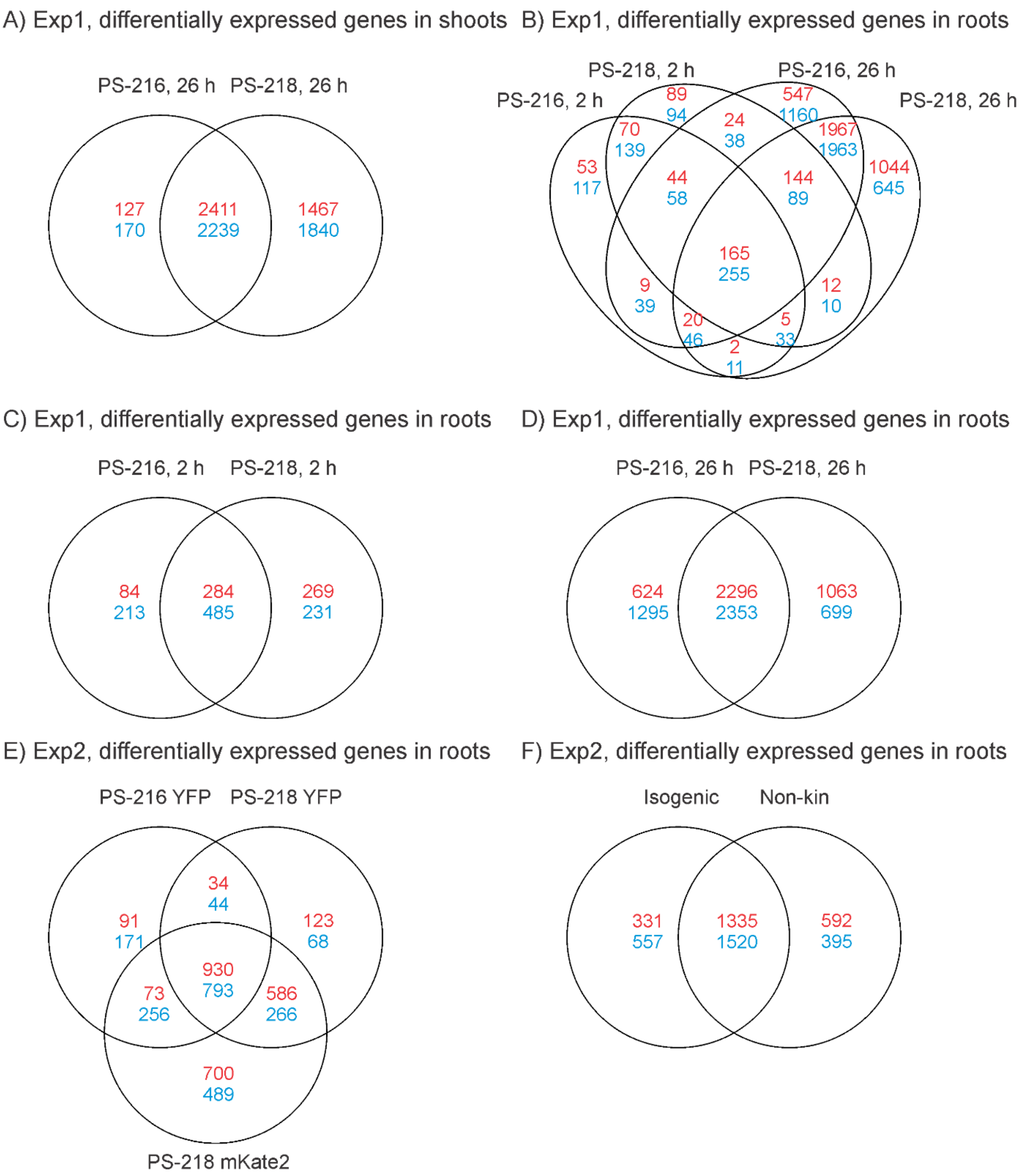


**Figure S3: Number of genes, regulated by *B. subtilis* in plants incubated in *B. subtilis* culture of high density ($10^7$ CFU/mL).** A) Shoots of the first experiment comparing strains PS-216 and PS-218 after 26-h incubation. B) Roots of the first experiment comparing strains PS-216 and PS-218 after 2-h and 26-h incubation. C) Roots of the first experiment comparing strains PS-216 and PS-218 after 2-h incubation. D) Roots of the first experiment comparing strains PS-216 and PS-218 after 26-h incubation. E) Roots of the second experiment comparing strains PS-216, PS-218 and PS-218 mKate2 after 2-h incubation. F) Roots of the second experiment comparing strains in isogenic and non-kin interactions after 2-h incubation. PS-216: PS-216 YFP, PS-218: PS-218 mKate2, isogenic: PS-218 YFP + PS-218 mKate2, non-kin: PS-216 YFP + PS-218 mKate2. Blue: downregulated genes, red: upregulated genes. In all Venn diagrams, only genes with absolute log2 fold change above 1 and FDR-adjusted p-values below 0.05 are counted. See Table S3 for results on individual genes.

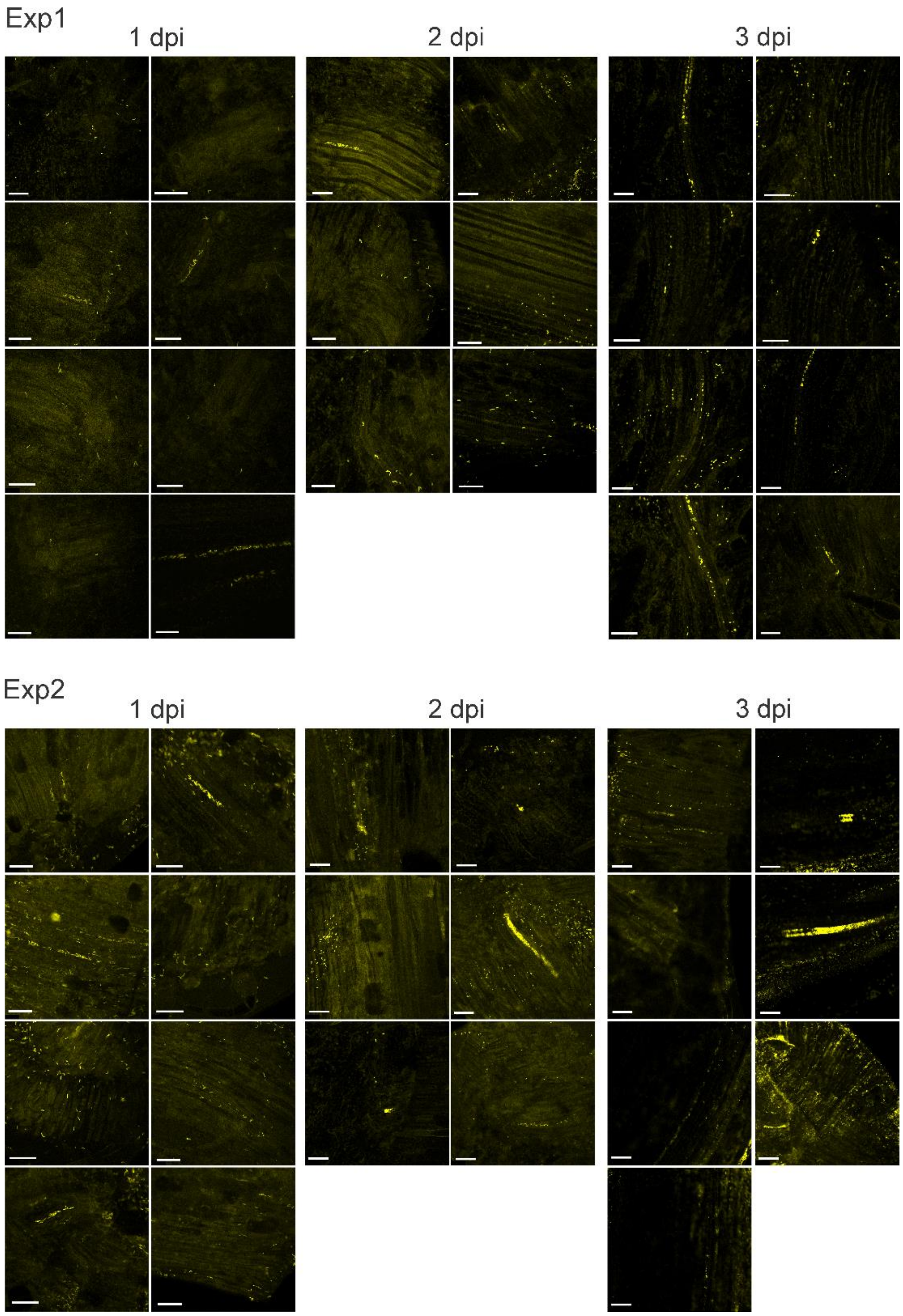


**Figure S4: Observation of *B. subtilis* cells in potato tissue after internalization.** Cells in potato cv. Rywal tissue were imaged by following YFP fluorescence up to three days after inoculation (dpi) of the tissue culture plants with *B. subtilis* PS-218 YFP culture. YFP fluorescence is coloured as yellow. Scale is 50 µm. Exp – experiment.

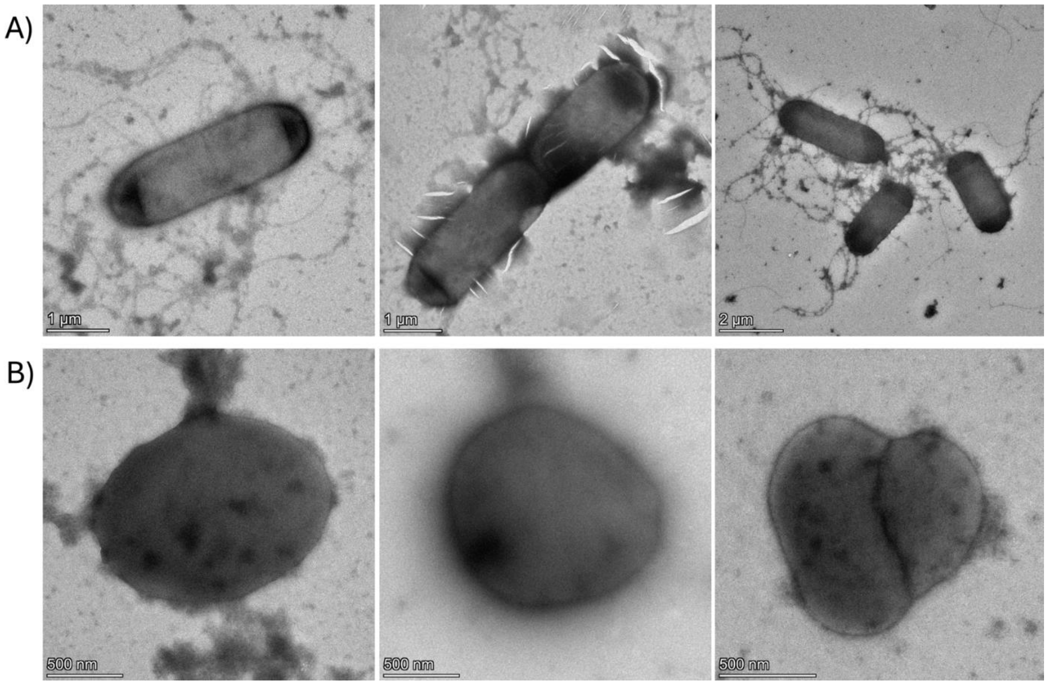


**Figure S5:** ***B. subtilis*** **cells as observed in liquid culture (A) and potato leaf homogenate (B).**

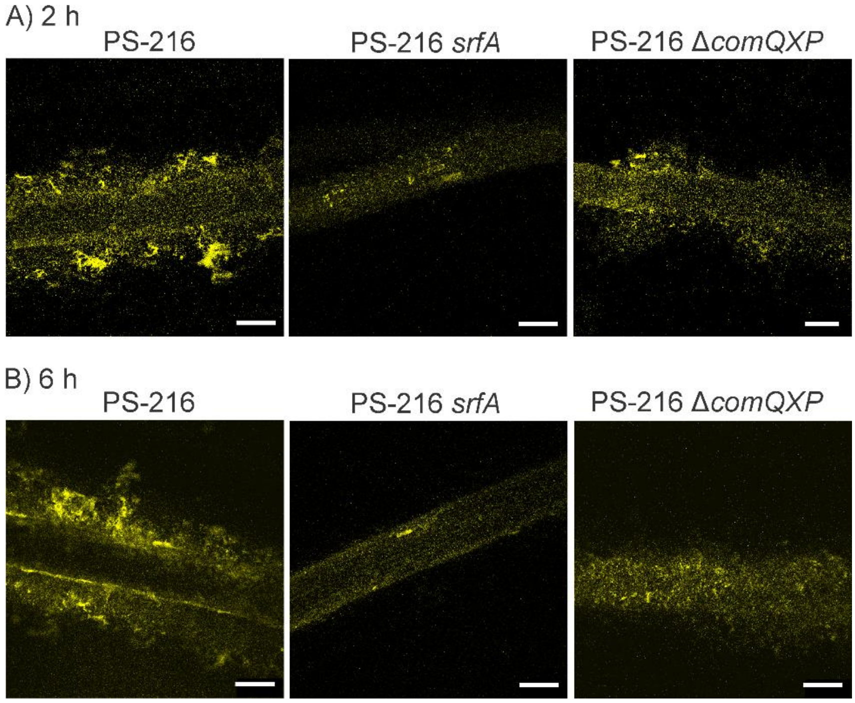


**Figure S6: Biofilm formation on potato roots inoculated with *B. subtilis* mutants that have attenuated surfactin production.** Biofilm formation was imaged on potato roots of cv. Rywal by confocal microscopy after A) 2 h and B) 6 h incubation in YFP-labelled *B. subtilis* PS-216, *B. subtilis* PS-216 *srfA* and PS-216 *ΔcomQXP* culture of high density ($10^7$ CFU/mL). YFP fluorescence is shown. Scale is 250 µm.

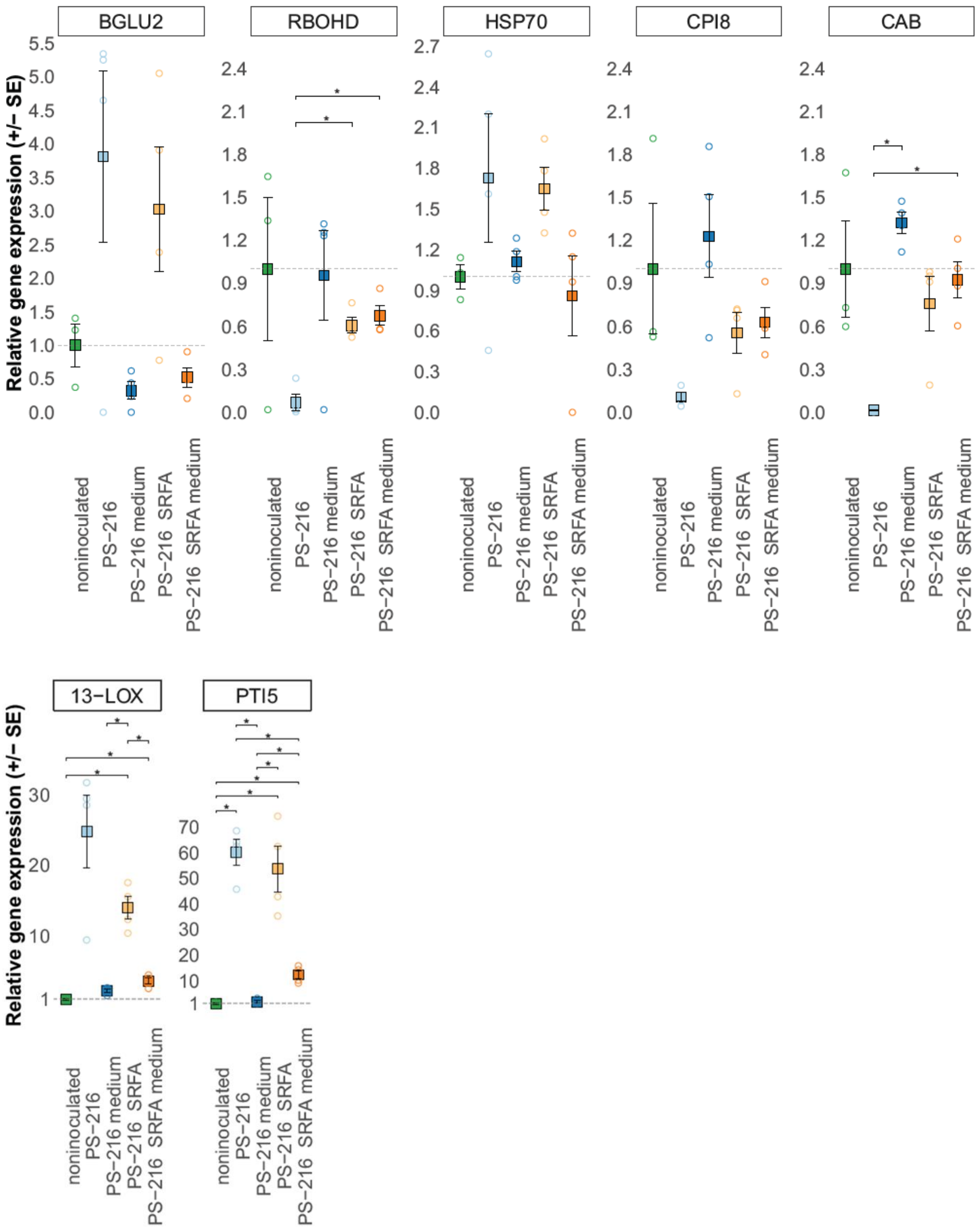


**Figure S7: *B. subtilis*-produced secondary metabolites in the medium do not contribute significantly to the PTI5 and 13-LOX upregulation.** Potato plants of cv. Rywal from tissue cultures were added to bacteria culture (YFP-labelled *B. subtilis* PS-216 and *B. subtilis* mutant PS-216 *srfA* with attenuated surfactin production) with OD600 0.02 and incubated overnight. After 17 h, plants and bacteria were discarded and new plants were added to conditioned medium and incubated for 26 h. As a positive control, plants were inoculated with YFP-labelled *B. subtilis* PS-216 and *B. subtilis* mutant PS-216 *srfA*

with attenuated surfactin production as in previous experiments. As a negative control, non-inoculated samples (inoculated with MS30 medium) were used. Relative expressions (relative to the endogenous control, see methods) of several genes (BGLU2, HSP70, RBOHD, 13-LOX, CAB, CPI8, PTI5) were measured in roots by quantitative PCR (qPCR). Relative gene expression was scaled to the average gene expression of non-inoculated group (marked with dotted line). Welch's t-test was used to determine differences between treatments. Mean and standard error of the mean are shown. Asterisks (*) denote statistically significant difference ($p$-value < 0.05).

isogenic

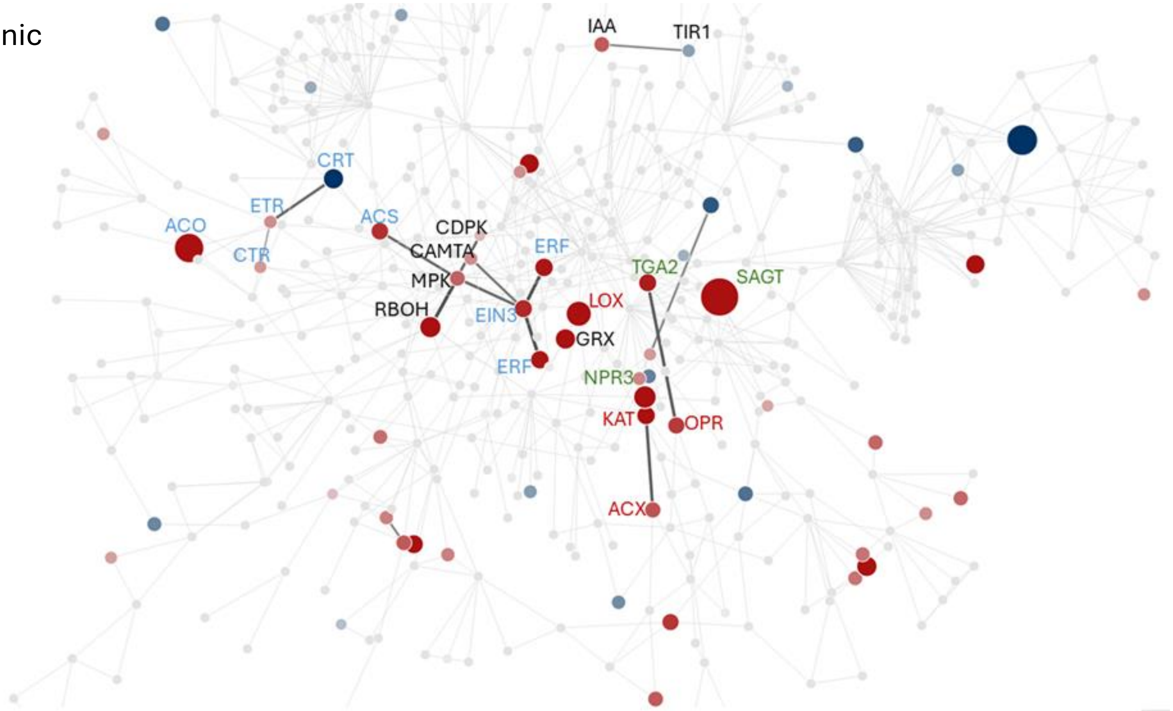


non-kin

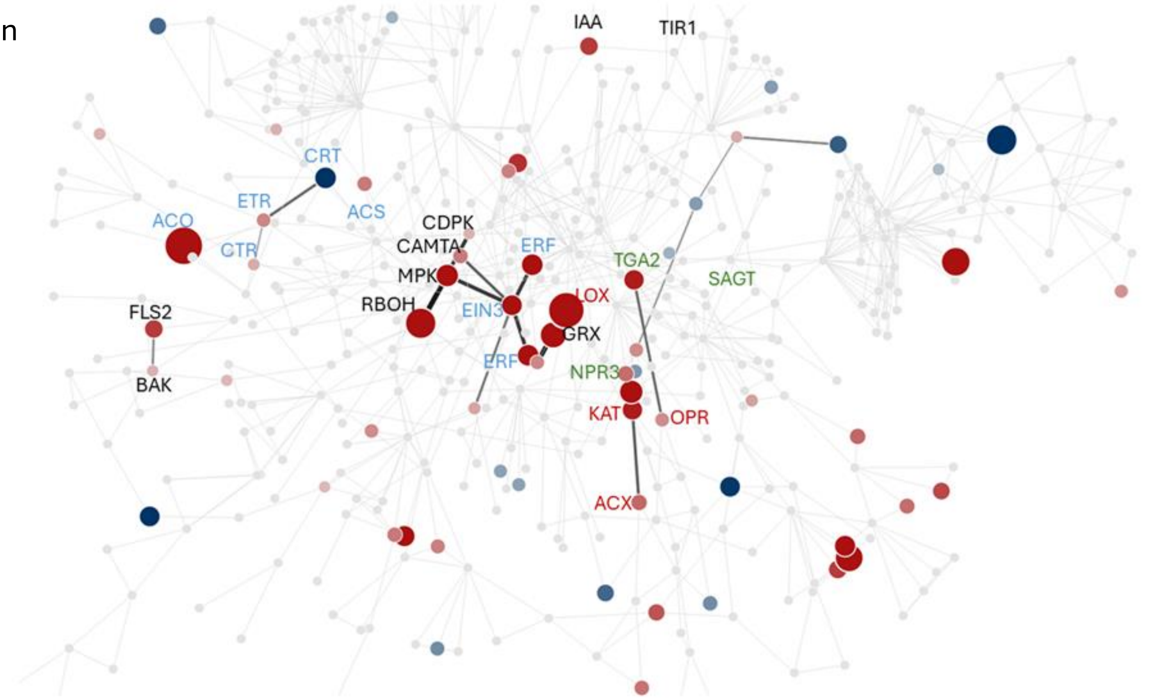


**Figure S8: Regulated central signaling modules in potato roots in isogenic and non-kin interactions.** Isogenic: PS-218 YFP + PS-218 mKate2, non-kin: PS-216 YFP + PS-218 mKate2. RNA-seq data was overlayed with plant stress signaling prior knowledge network, built from experimental data on protein-protein interactions, protein-DNA interactions, and metabolic pathways (skm.nib.si)[35] to determine regulated central signaling modules in isogenic and non-kin interactions. Genes from ethylene (blue), JA (red), SA (green) and other (black) signaling modules are shown. See Table S7 for the name and description of regulated genes.

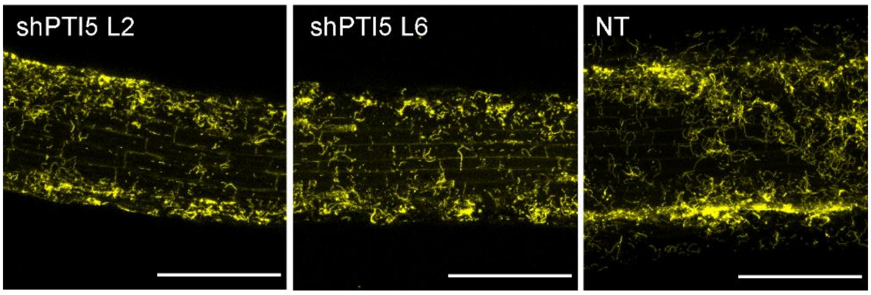


**Figure S9:** ***B. subtilis*** **root colonization of transgenic potato lines with silenced PTI5 (shPTI5 L2 and L6) and non-transgenic (NT) potato plant.** Plants were added to *B. subtilis* PS-218 YFP culture of high density ($10^7$ CFU/mL), incubated for 2 h and imaged under confocal microscope. YFP fluorescence is coloured as yellow. Scale is 250 µm.

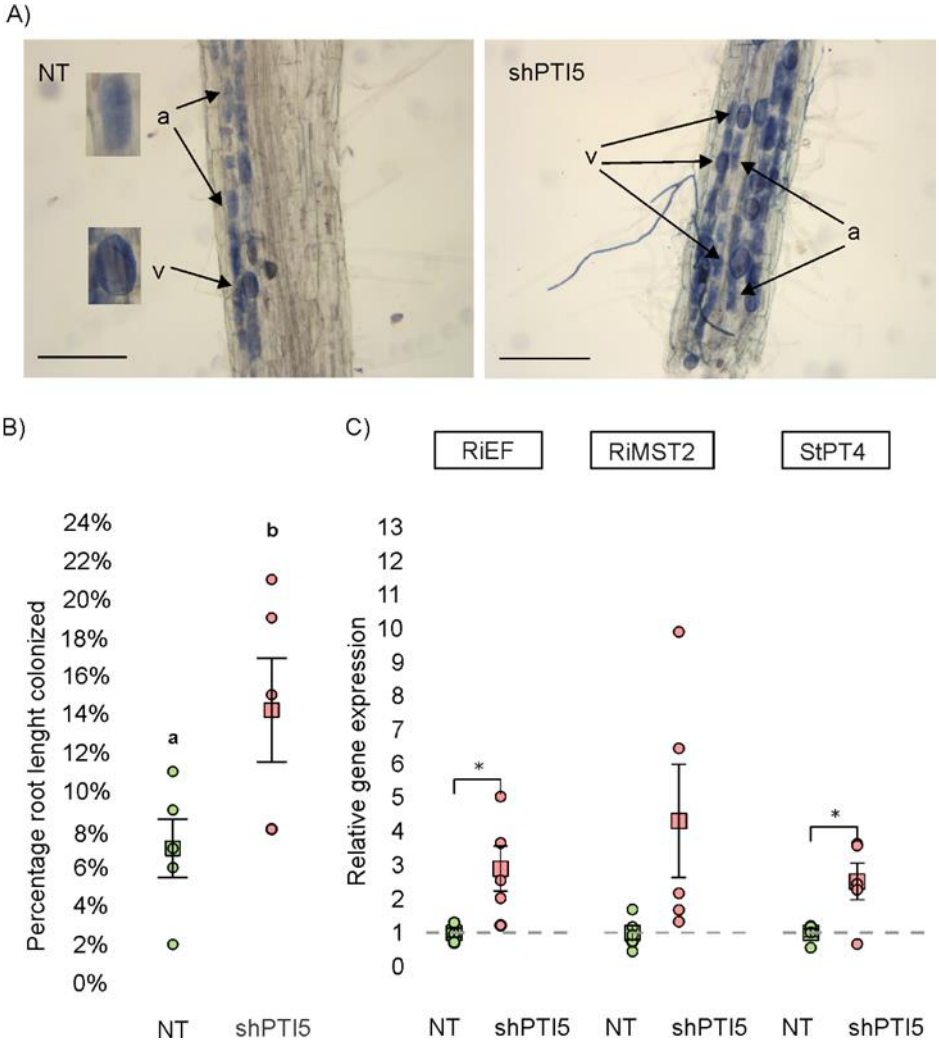


**Figure S10: Mycorrhizal colonization of potato roots in non-transgenic potato plant (NT) and PTI5-silenced genotype line 2 (shPTI5).** A) Representative images of mycorrhizal colonization in the different genotypes. Fungal structures within the root cortex of potato plants are stained in blue (a: arbuscules, v: vesicles). Scale is 200 µm. B) Percentage of root length colonized by the arbuscular mycorrhizal fungi *R. irregularis* C) Expression analysis of *R. irregularis* constitutive gene (RiEF) and symbiosis marker genes, the *R. irregularis* monosacaride transporter (RiMST2) and the potato phosphate transporter (StPT4), as well as regression with Dunn's pair-wise comparisons were used to determine the differences between the genotypes. Different letters show statistical differences. Permutation test was used to determine the differences between the genotypes. Individual measurements (circles), mean (squares) and standard error are shown. Asterisks denote a statistically significant difference (*: p-value < 0.05, **: p-value < 0.01).
